# Ankyrin domain encoding genes resulting from an ancient horizontal transfer are functionally integrated into developmental gene regulatory networks in the wasp *Nasonia*

**DOI:** 10.1101/383968

**Authors:** Daniel Pers, Jeremy A. Lynch

## Abstract

**Background:** How and why regulatory networks incorporate additional components, and how novel genes are maintained and functionally integrated into developmental processes are two important and intertwined questions whose answers have major implications for the evolution of development. We recently described a set of novel genes with robust and unique expression patterns along the dorsal-ventral axis of the embryo of the wasp *Nasonia.* Given the unique evolutionary history of these genes, and their apparent integration in to the dorsal-ventral (DV) patterning network, they are collectively an excellent model to study the evolution of regulatory networks, and the fates of novel genes.

**Results:** We have found that the novel DV genes are part of a large family of rapidly duplicating and diverging ankyrin domain encoding genes that originated most likely by horizontal transfer from *Wolbachia* in a common ancestor of the wasp superfamilly Chalcidoidea. We tested the function of those ankyrin encoding genes expressed along the DV axis and found that they participate in early embryonic DV patterning. We also developed a new wasp model system *(Melittobia)* and found that some functional integration of ankyrin genes have been preserved for over 90 million years, while others are lineage specific.

**Conclusions:** Our results indicate that regulatory networks can incorporate novel genes that then become necessary for stable and repeatable outputs. Even modest role in developmental networks may be enough to allow novel or duplicate genes to be maintained in the genome and become fully integrated network components.

## Background

Gene regulatory networks (GRNs) coordinate the expression of mRNA and proteins in a spatiotemporal manner to bring about a specific developmental output [1. The complex webs of interacting nodes and modules that make up GRNs are vital for establishing patterning, morphogenesis, and ultimately an organism’s body plan [2]. Perturbations to these networks should result in novel developmental outputs. However, canalization and developmental redundancy can conceal underlying genetic variation and phenotypic plasticity. This quality enables GRNs to weather large variations in genomic and environmental inputs, without disrupting the phenotypic output of the network [3-6].

These properties of GRNs raise questions about how developmental mechanisms can evolve. Since robust networks can absorb large genetic changes without causing major changes in developmental output, it seems that a large threshold must be overcome in order to achieve a new phenotype [7]. Thus, robustness paradoxically may make GRNs less able to respond to evolutionary pressures, since most mutations will not produce phenotypes visible to natural selection. Thus, we might expect robust developmental GRNs to be mostly static in evolutionary time in the absence of major phenotypic change. However, there are many well known examples where developmental processes have been apparently unchanged, while the molecular basis of development is highly diverged [8-10]

Whether these changes are fixed because they provide some selectable improvement on the developmental process of interest, are indirect responses to selection on modules that are reused in other developmental processes, or are random is not well characterized. The development of methods to circumvent the candidate gene approach in a very wide variety of species facilitates comprehensive characterization of developmental GRNs at high phylogenetic resolution. This can allow hypotheses about the evolution of development to be tested robustly, and will lead to a deep understanding of the pattern and process of GRN evolution.

The GRN that patterns the embryonic dorsoventral (DV) axis of the wasp *Nasonia vitripennis* has been shown to be a good model to study novelty and the evolution of gene networks. Having split from *Drosophila melanogaster* over 300 MYA [11], the two have converged on a similar mode of embryogenesis [12], and share a nearly identical expression of tissue-specific marker genes just prior to gastrulation [13]. We have previously shown that most genes differentially expressed along the DV axis of the *Nasonia* embryo are not conserved components of the *Drosophila* DV GRN, making the comparison between the fly and wasp DV GRNs an ideal system for understanding how GRNs evolve while producing similar patterning results [14, 15].

A particularly interesting case of *Nasonia* specific DV GRN components are a set of 15 ankyrin domain containing genes, which do not have clear orthologs in *Drosophila* or in any other insects outside of the Superfamily Chalcidoidea. In fact, there is evidence that these genes entered the genome of the ancestor of *Nasonia* through at least one horizontal gene transfer (HGT) event, followed by several waves of duplication and divergence. We have previously shown that these ankyrin domain encoding genes are expressed in specific patterns along the DV axis [15], and here we demonstrate that they also are functionally incorporated into the DV patterning GRN, and their loss leads to variable disruptions to patterning. Through examination of another wasp, *Melittobia digitata,* we also show that some of the functional incorporation is ancient within the Superfamily, while there is also strong evidence of recent gains and/or losses of function in the *Nasonia* and *Melittobia* lineages.

We propose that the properties of ankyrin domain containing proteins allow them to rapidly gain interaction partners, and potentially adaptive functions in developmental networks, which increases the likelihood that genes of this type will be maintained and sometimes multiply in the course of genome evolution.

## Results

### Identification of new families of apparently horizontally transferred, ankyrin domain encoding, genes

In our previous study we identified fifteen transcripts encoding ankyrin domain proteins that appeared to be significantly regulated by the Toll and/or BMP signaling pathways in the *Nasonia* embryo [15]. Further analysis of their expression showed that 6 of these genes are expressed laterally, 3 are expressed on the dorsal surface of the embryo, one is expressed over the ventral midline, one has a complex pattern involving late expression in dorsal tissues, and 4 with no clear differential expression along the DV axis (described in more detail below).

Our previous analysis indicated that four of these 15 genes possess a PRANC (**<u>P</u>**ox proteins **R**epeats of **<u>AN</u>**kyrin, **<u>C</u>**-terminal) domain at their C-termini. PRANC domains were originally described in Pox viruses. They were first described in a eukaryotic system upon the publication of the *Nasonia* genome, where a set of PRANC domain encoding genes were found to be integrated into the genome, and which are highly similar PRANC domain proteins in their endosymbiotic bacteria, *Wolbachia*[16, 17]. The similarity (both in the PRANC domains, and their association with ankyrin repeats) of the PRANC encoding genes integrated into the *Nasonia* genome to those found in the *Wolbachia* genome led to the hypothesis that the *Nasonia* PRANC genes originated from a HGT from *Wolbachia*.

While the remaining 11 DV regulated ankyrin domain encoding genes do not appear to have PRANC domains, we believe that they entered the *Nasonia* genome through similar processes of horizontal transfer, gene duplication and rapid molecular divergence.

If we focus on BLASTp searches using any of the 15 *Nasonia* DV ankyrin domain genes as queries against well annotated genomes from *D. melanogaster* or *Apis meliferra* the results are invariably “canonical” ankyrin genes (i.e., *ankyrin-1* or *ankyrin-* 2), that are highly conserved in all insect species. As expected, *Nasonia* possesses clear direct orthologs to such canonical ankyrin genes that are highly conserved at the amino acid level and these gene families are clearly distinct from the genes we propose are horizontally transferred (Additional File 1 and 2).

Additional evidence that these genes originated outside of the normal course of vertical transmission from generation to generation is the pattern of unrestricted BLASTp results and the phylogenetic relationships of the top 100 protein BLAST hits for each *Nasonia* ankyrin (Additional File 3) which do not follow expected phylogenetic patterns (i.e., stronger hits throughout the hymenoptera, then hits in Diptera, Coleoptera, Lepidoptera, other insects, etc…), that are observed for the canonical ankyrin genes.

As would be expected, the top hits for most of the 15 *Nasonia* sequences come from *Trichomalopsis* (a sister genus to *Nasonia),* except one where there is a recent *Nasonia* paralog almost identical to it (Additional File 3). This indicates that the ancestors of the 15 DV regulated ankyrin genes were present in the common ancestor of the *Nasonia* and *Trichomalopsis* genera. In addition, sequences from the fig wasp *Ceratosolen solmsi* [18], Copidosoma floridanum and *Trichogramma pretiosum* [19] (representing the Families Agaonidea, Encyrtidae, and Trichogrammitidae within the Superfamily Chalcidoidea, respectively) are among the top hits for most of the *Nasonia* proteins, and cluster near to the *Nasonia* query sequence in the phylogenies (Additional File 3).

Outside of the chalcid wasps, strong hits are scattered across the tree of life. Taxa with proteins showing strong similarity to each *Nasonia* DV ankyrin gene include the ant *Pseudomyrmex gracilis,* the single celled eukaryote *Trichomonas vaginalis,* the bee *Ceratina capitata,* the sea urchin *Strongylocentrotus purpuratus,* and the amoeba endosymbiont *Candidatus Amoebophilus asiaticus* [20]. Other taxa occur with less frequency, including *Wolbachia* sequences (Additional File 3). Importantly, canonical ankyrins from *Nasonia* or other insect species do not appear among the top hundreds of hits and do not cluster with our genes of interest in phylogenies.

We attempted to phylogenetically analyze the Nasonia DV-ankyrin genes in the context of their best blast hits in other animals and microbes to see if they cluster with prokaryotic or canonical eukaryotic ankyrin sequences. However, we found that aligning our relatively small, rapidly evolving proteins with proteins encoding varying numbers of ankyrin domains of varying conservation made phylogenetic analyses difficult, and in our hands uninformative (not shown).

Instead, we decided to focus on the regions C-terminal to the ankyrin repeats in the *Nasonia* DV ankyrin domain genes. These regions range from ~100-200 amino acids, except in two sequences (-D and -E), which lack C-terminal sequence beyond the ankyrin domains. We knew that this region was predicted to contain a PRANC domain in four of our DV ankyrin proteins, but we did not know how similar to sequences outside of the wasps it would be. We also hypothesized that the remaining C-termini maintained cryptic similarity to the ancestral PRANC domain.

Because the C-termini of our DV ankyrin domain genes seem to be less well conserved, we used the more sensitive, iterative PSI-BLAST [21] approach to identify similar sequences in the NCBI nonredundant (nr) database. We used default parameters, including only using genes that were above the threshold in round one as templates to generate the pattern for the second-round search. We then took all of the aligning sequences that were above the significance threshold given by PSI-BLAST and subjected them to phylogenetic analysis. We only discuss genes that were above the threshold in the second round (with the exception of Nv-CLANK-L, which required 4 rounds), as this was sufficient to identify the first non-insect and microbial sequences. DV ankyrins -D, and -E were not analyzed as they lack sequence C-terminal to the ankyrin domains.

The taxa that appear in these queries are much more limited than what was found using the full DV ankyrin protein sequences as queries, indicating that complexity of aligning the constrained and repetitive ankyrin domains can indeed give spurious signals of homology. The close relationship of *Nasonia* DV ankyrins and other orphan ankyrin domain encoding genes in Chalcidoidea is accentuated, as multiple sequences from *Ceratosolen, Copidosoma,* and *Trichogramma* cluster more robustly with large numbers of *Nasonia* sequences in this analysis (Figure 1A, for example, Additional File 4). We also find sequences from species that also showed up strongly when using the full protein as a query, particularly from the ant *Pseudomyrmex* and bee *Ceratina*. We also find a large number of hits from the Braconid wasp *Microplitus demolitor* and the whitefly *Bemisia tabaci* in all PSI-BLASTs. The sequences from these insects cluster closely with others from the same species, and also with particular *Wolbachia* sequences (Figure 1B, for example, Additional File 4), again indicating recent HGT in these organisms.

**Fig 1.**
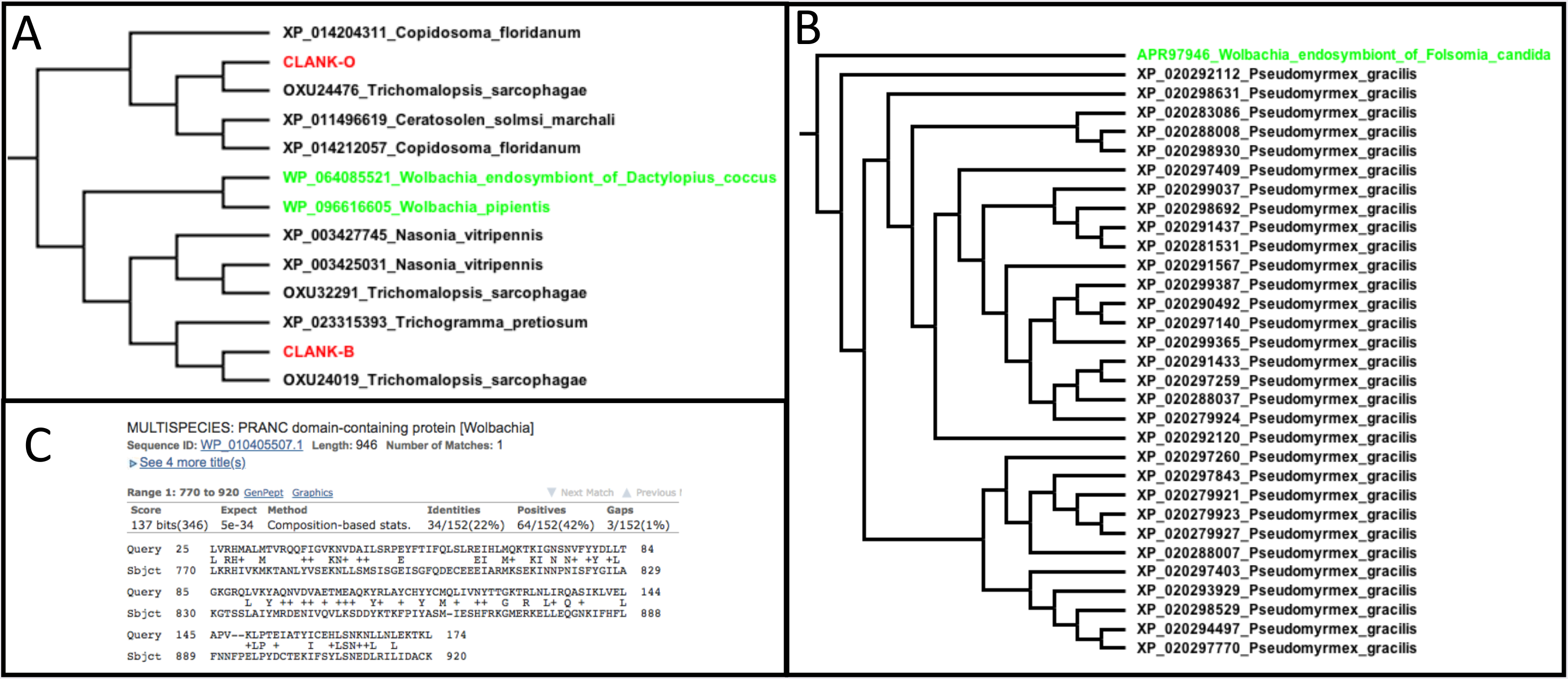
PSI-BLAST and phylogenetic analyses support the HGT origin of DV-regulated ankyrin genes in chalcid wasps (CLANKs), and additional HGTs in insects. A. A portion of the phylogenetic tree generated from significant sequences using the C-terminus of *Nv-CLANK-C* as a query in PSI-BLAST. DV regulated ankyrin genes CLANK-O and -N cluster with ankyrin genes from *Wolbachia,* and other representatives of the Chalcidoidea *(Trichogramma, Ceratosolen,* and *Copidosoma)*. **B**. Numerous proteins from the ant Pseudomyrmex cluster strongly and consistently with other Wolbachia ankyrin domain encoding genes. **C**. Screen capture of BLAST result illustrating the strong similarity of C-terminal sequence from a protein not annotated as having a PRANC domain *(Nv-CLANK-B)*.

Outside of these insect hits were primarily from prokaryotes and some poxviruses. *Wolbachia* species had the strongest and most common prokaryotic hits, but many *Orientia* (a Rickettsial intracellular parasite where PRANC domains were also found in the *Nasonia* genome publication [17]) sequences were also found (Additional File 4). The e-values of Wolbachia hits ranged from E-05 to E-60 (Fig. 1C, for example). Importantly, all of the hits aligned to the C-terminal region of small *Wolbachia* proteins that also have ankyrin domains toward the N-terminus. Some of the hits in these analyses are annotated as PRANC domains, but the majority of them are not, despite highly significant similarity.

Overall, these results strongly indicate that the ankyrin domain containing genes that were identified in the PSI-BLAST analyses have a common history, despite the fact that they occur in a scattered phylogenetic distribution in a few insect lineages, intracellular bacteria, and viruses. We find it extremely unlikely that these proteins with conserved C-terminal, PRANC-like motifs that are directly downstream from relatively well conserved ankyrin domains, would have evolved convergently by chance multiple times.

Rather, we believe that the pattern we uncovered indicates multiple instances of HGT: At least four recent ones in lineages leading to *Pseudomyrmex, Bemisia, Ceratina*, and *Microplitus,* and one ancient one in a common ancestor of the superfamily Chalcidoide (around 150 million years ago [22]). We propose to name this later family of proteins Chalcidoidea Lineage specific **ANK**yrin domain encoding genes (***<u>CLANK</u>**s*). We will henceforth discuss the DV ankyrin domain proteins as *Nasonia vitripennis* CLANKs (Nv-CLANK) -A through -O. The relationships between our CLANK nomenclature and gene identification numbers in different annotations are given in Additional File 5.

While we strongly favor the hypothesis that the CLANKs entered the genomes of Chalcid wasps through HGT based on the evidence given above, the complex relationships and genetic exchange back and forth among prokaryotes, viruses and eukaryotes [16, 20, 23], make proving this idea beyond a shadow of a doubt a daunting task, well beyond the scope of this manuscript. Whatever the case, CLANKs are new genes in the wasps relative to the rest of the insects, and we would like to understand why they have been maintained over the course of more than 150 million years of evolution in this clade.

### Characterization of protein and genomic sequence features

Characterization of the amino acid sequences and genomic context of these CLANK genes supports that this family of genes likely has a long history, and has gone extensive sequence change in the course of their evolution. These 15 genes code for 18 proteins that vary in size from 255 to 717 amino acids. Most of the genes encoding *Nv-CLANKs* are interrupted by at least one intron, and they are found spread across the 5 chromosomes of *Nasonia* (Fig. 2). Sequence alignment of these 15 protein sequences yielded very few residues conserved across the CLANKs [24] (data not shown). Phylogenetic analysis [25] of the DV CLANKs gives two major clades (Fig. 2). Domain analysis with Interproscan [26] confirmed the four CLANKS have officially annotated PRANC domains (Nv-CLANK-F, -J, -N, -O), and mapping these on the phylogeny shows that this domain has likely been lost or degraded beyond recognition in many lineages (Fig. 2). This analysis also showed that the number of Ankyrin repeats within a protein varies from 6 to 13 throughout the length of the proteins, except for the last ~100 amino acids, where the PRANC-like domains reside.

**Fig 2.**
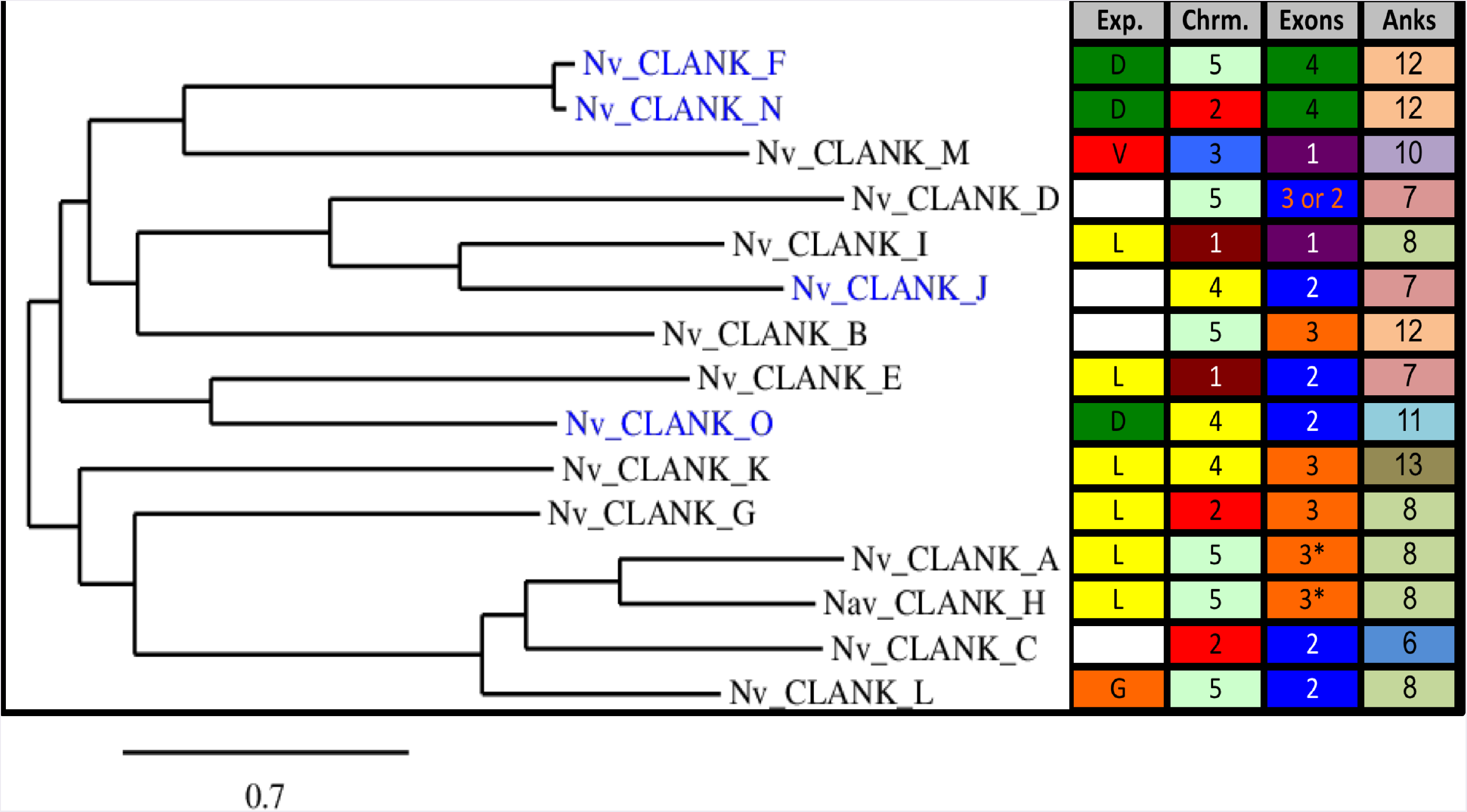
Phylogenetic analysis of *Nasonia* DV regulated CLANKs protein family. (Left) Phylogenetic tree of CLANK proteins of interest generated using “One Click” Phylogeny Analysis (http://phylogeny.lirmm.fr) [25]. Branch length is proportional to the number of substitutions per site. Blue text represents proteins containing a PRANC domain. (Right) Corresponding mRNA expression domain (D = dorsally, L = laterally, V = ventrally, G = no diff. expression until gastrulation, blank = ubiquitous or lack of expression), chromosomal location, number of exons (* = two differentially spliced 43 transcripts occur, both contain same number of exons), and number of ankyrin n repeats for each CLANK protein of interest. Colors added to emphasize similar value in each column.

Overall, there appears to be no correlations between these genetic features and regulation, expression patterns, or phylogenetic relatedness (Fig. 2), except for the two pairs of recent paralogy (Nv-CLANK-F and -N, and -A and -H).

### Detailed characterization of DV CLANK embryonic expression

RNA expression patterns for the 15 CLANK genes have previously been mentioned [15], but not fully described. Thus, *in situ* hybridization experiments were thus repeated and analyzed in more detail over a longer developmental time frame, and transcripts were grouped according to their expression patterns. Four CLANKS *(-B, -C, -D, -J)* have no patterned expression at any time in embryogenesis (Additional File 6) and will not be discussed further.

### Laterally expressed CLANKs

Six of the fifteen *Nv-CLANK* transcripts are expressed in a lateral domain at one or more time points during embryogenesis; however, this expression is quite dynamic. Three *Nv-CLANKs* (-G, -H, -K) show an overall expansion of expression, while three *Nv-CLANKs* (*-A, -E, -I*) are characterized by a shrinking of their expression domain.

*Nv-CLANK-G* is ubiquitously expressed during the pre-blastoderm and early blastoderm stages of development (Fig. 3A1-32). As the blastoderm undergoes additional rounds of division and begins to cellularize, *Nv-CLANK-G* is expressed first as a band encircling the anterior end of the embryo (Fig. 3A3) and then expands posteriorly creating a gradient with highest levels in that initial anterior domain (Fig. 3A4). Expansion is missing from both poles, and also from the dorsal midline of the embryo (Fig. 3A5). During gastrulation, expression is restricted to a segmental pattern in the cells that give rise to the central nervous system (CNS) (Fig. 3A6).

**Fig 3.**
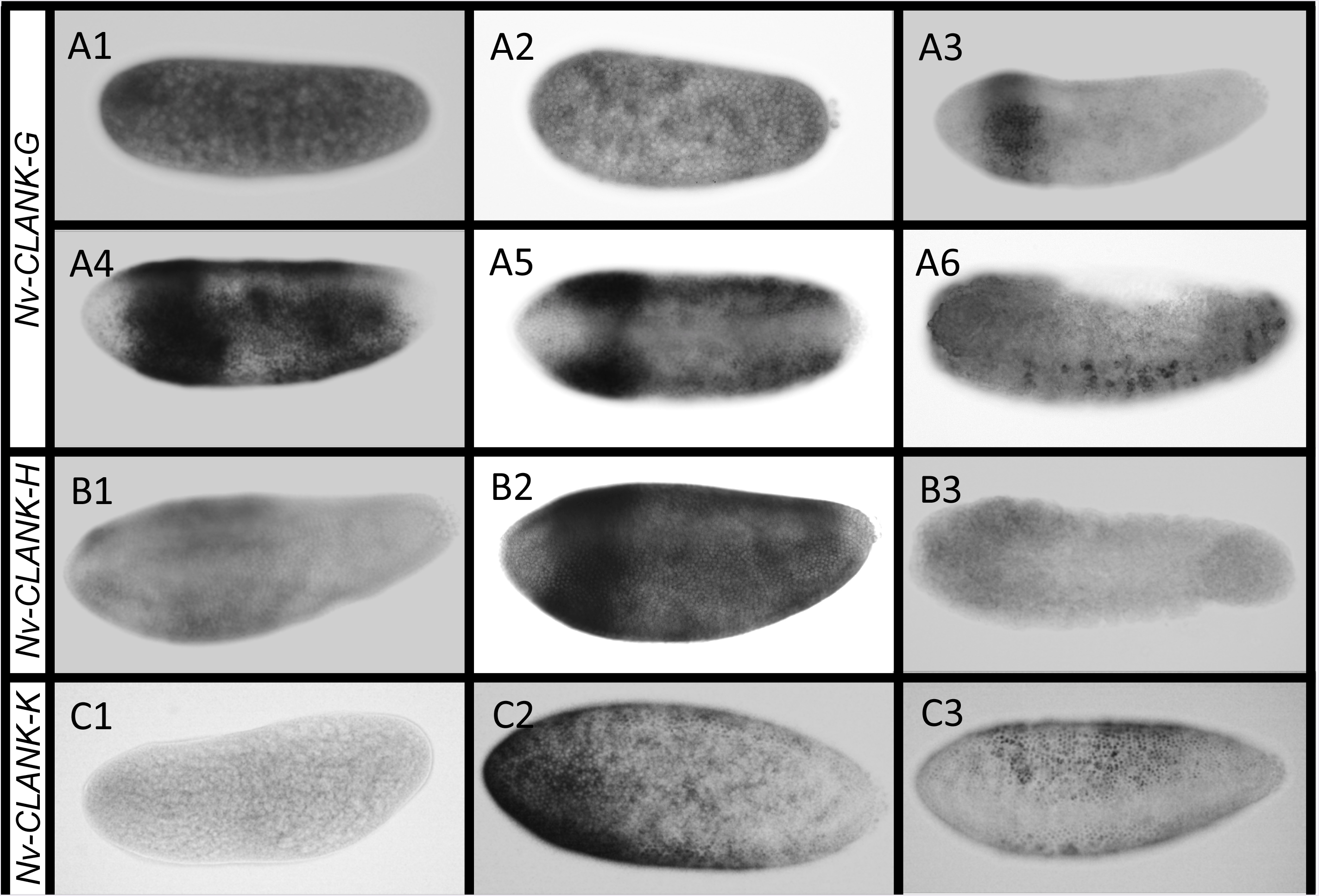
Laterally expressed CLANKs with dynamic expansion. A1-A6. Expression of *Nv-CLANK-G* from pre-blastoderm through gastrulation. **B1-B3** Expression of *Nv-CLANK-H* from blastoderm to gastrulation. **C1-C3** Expression of *Nv-CLANK-K* from pre-blastoderm through late-blastoderm. All embryos are oriented with anterior to the left, posterior to the right, dorsal up, and ventral down (except A5, dorsal view).

The expression pattern of *Nv-CLANK-H* is very similar to *Nv-CLANK-G* during the middle and late blastoderm stages. It also is initially expressed in an anterior band that eventually forms a gradient of expression along the AP axis (Fig. 3B1-2). Again, expression is absent at the AP poles and along the dorsal midline. However, while *Nv-CLANK-G* was expressed maternally, *Nv-CLANK-H* has no early expression (data not shown) and is ubiquitously expressed at very low levels during gastrulation instead of being localized to CNS precursors (Fig. 3B3).

*Nv-CLANK-K* is expressed maternally and ubiquitously at low levels (Fig. 3C1) before being expressed in an anterior-lateral domain with expression lacking at both the ventral and dorsal midline and strongest near the anterior pole (Fig. 3C2). This domain then shifts and expands posteriorly into a more evenly expressed lateral domain, with inhibition of expression ventrally and at the poles (Fig. 3C3). During gastrulation expression is lost completely (data not shown).

*Nv-CLANK-A* is initially expressed ubiquitously at very low levels before becoming localized in a broad lateral domain in the early blastoderm (Fig. 4A1-A2). The lateral domain then shrinks into two discrete bands in the trunk of the embryo, before expanding first dorsoventrally and then anteroposteriorly into one lateral band (Fig. 4A3-A5). At times, expression is missing along the ventral and dorsal midlines; however, this is variable and dynamically changes as the blastoderm undergoes further division and cellularizes. Expression is lacking in the gastrulating embryo (Fig. 4A6).

**Fig 4.**
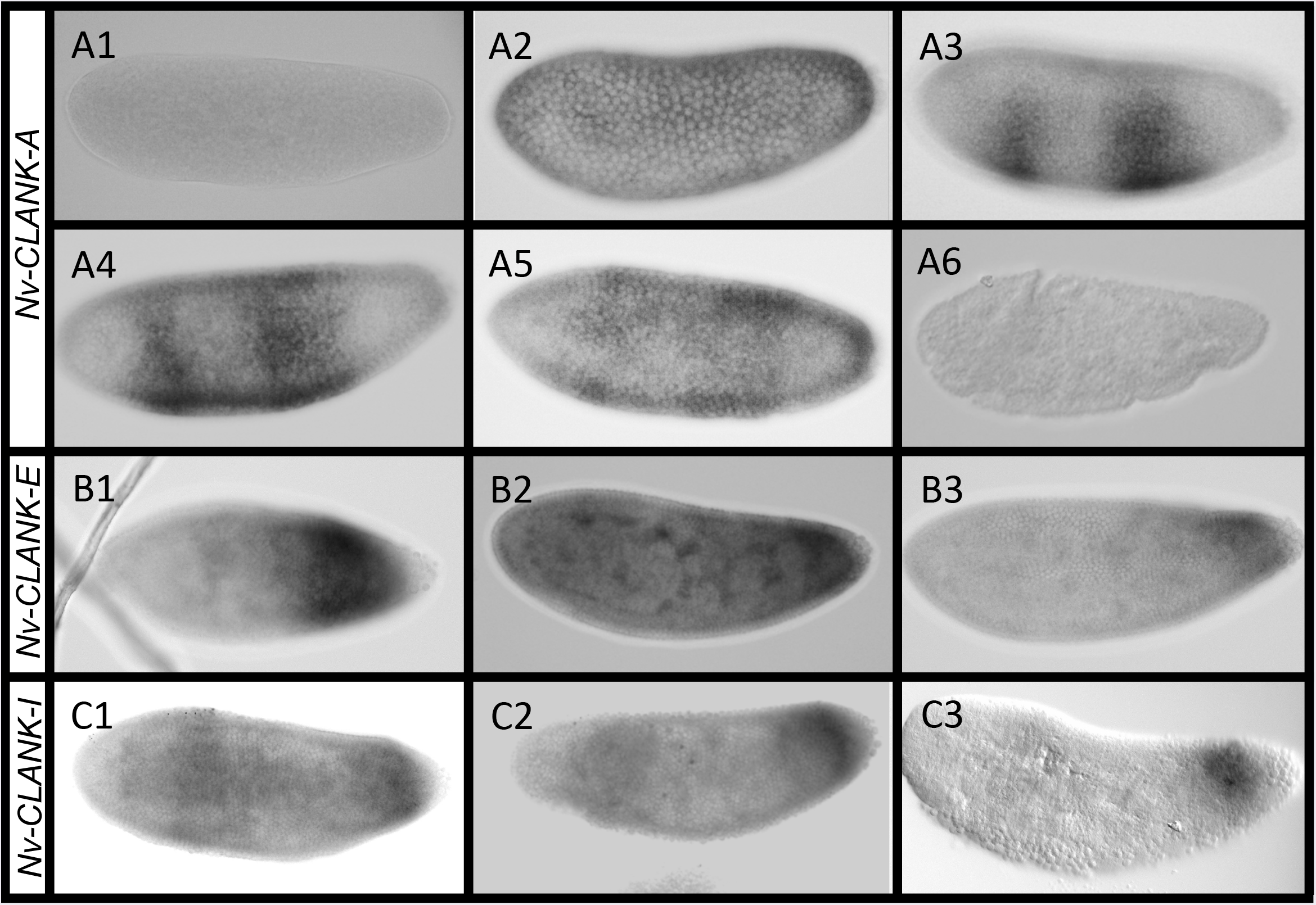
Laterally expressed CLANKs with dynamic retraction. A1-A6. Expression of *Nv-CLANK-A* from pre-blastoderm through gastrulation. **B1-B3** Expression of *Nv-CLANK-E* from mid to late blastoderm. **C1-C3** Expression of *Nv-CLANK-/* from blastoderm through gastrulation. All embryos are oriented with anterior to the left, posterior to the right, dorsal up, and ventral down.

*Nv-CLANK-E* expression is lacking or at very low levels both before and after the blastoderm stages (data not shown). Expression is in a broad band from the nearly the middle of the embryo to just anterior of the posterior pole during the syncytial blastoderm (Fig. 4B1). During cellularization, this band retracts to a smaller, weaker, expression domain in the posterior of the embryo (Fig. 4B2-B3).

The expression pattern of *Nv-CLANK-/* in the mid to late blastoderm also resembles that of *Nv-CLANK-G* and *-H*. Again, there is a lateral expression domain with expression missing from the two poles and the dorsal midline (Fig. 4C1); however, the dynamics of this transcript differ. Instead of expression first appearing in the anterior region and then appearing progressively towards the posterior pole, *Nv-CLANK-/* is initially expressed in this broad domain, spanning the trunk of the embryo, having slightly higher expression in the anterior and posterior regions. During gastrulation, the anterior expression is lost until there is just a posterior expression band (Fig. 4C2) and then a posterior spot (Fig. 4C3). *Nv-CLANK-/* is not expressed maternally or in the early blastoderm. (data not shown).

### Dorsally expressed CLANKs

Three of the fifteen transcripts are expressed dorsally during embryogenesis. *Nv-CLANK-N* and *-O* both lack maternal expression and are first expressed in the syncytial blastoderm. *Nv-CLANK-N* is expressed strongly at the anterior and posterior poles and appears to have weak and variable expression along the dorsal midline (Fig. 5A1-A3). *Nv-CLANK-O* is expressed evenly along the dorsal midline from pole to pole, stably expressed throughout syncytial divisions and cellularization (Fig. 5B1-B2). While *Nv-CLANK-N* lacks expression during gastrulation, *Nv-CLANK-O* is expressed dorsally, surrounding the extraembryonic material (Fig. 5B3).

*Nv-CLANK-F* is initially expressed at a low ubiquitous level (Fig. 5C1) before gaining expression at the anterior and posterior poles (Fig. 5C2). Expression then expands along the dorsal midline, anterior to posteriorly, ultimately connecting the two poles (Fig. 5C2-C4). In addition, there is dynamic ventral expansion that varies from embryo to embryo and from stage to stage. In some cases, the dorsal stripe expands past the anterior pole, onto the ventral half of the embryo, creating an anterior cap (Fig. 5C4). This expansion is accompanied by a perpendicular band, encircling the posterior end of the embryo’s trunk (Fig. 5C5). In other cases, the expansion does not cross into the ventral half, but still expands to broad domains at the two poles, while remaining narrow in the mid-trunk region (Fig. 5C6). As the onset of gastrulation nears, this expansion, and the initial dorsal stripe, retracts, leaving a strong patch of expression at the anterior pole, a lighter and smaller domain at the posterior pole, and a weak stripe perpendicular to the dorsal midline in the anterior-dorsal region of the embryo (Fig. 5C7). While the anterior patch remains strong at the start of gastrulation, the other two expression domains weaken quickly (Fig. 5C8). Differential expression is eventually lost during gastrulation, and the whole embryo exhibits weak, ubiquitous expression (Fig. 5C9).

**Fig 5.**
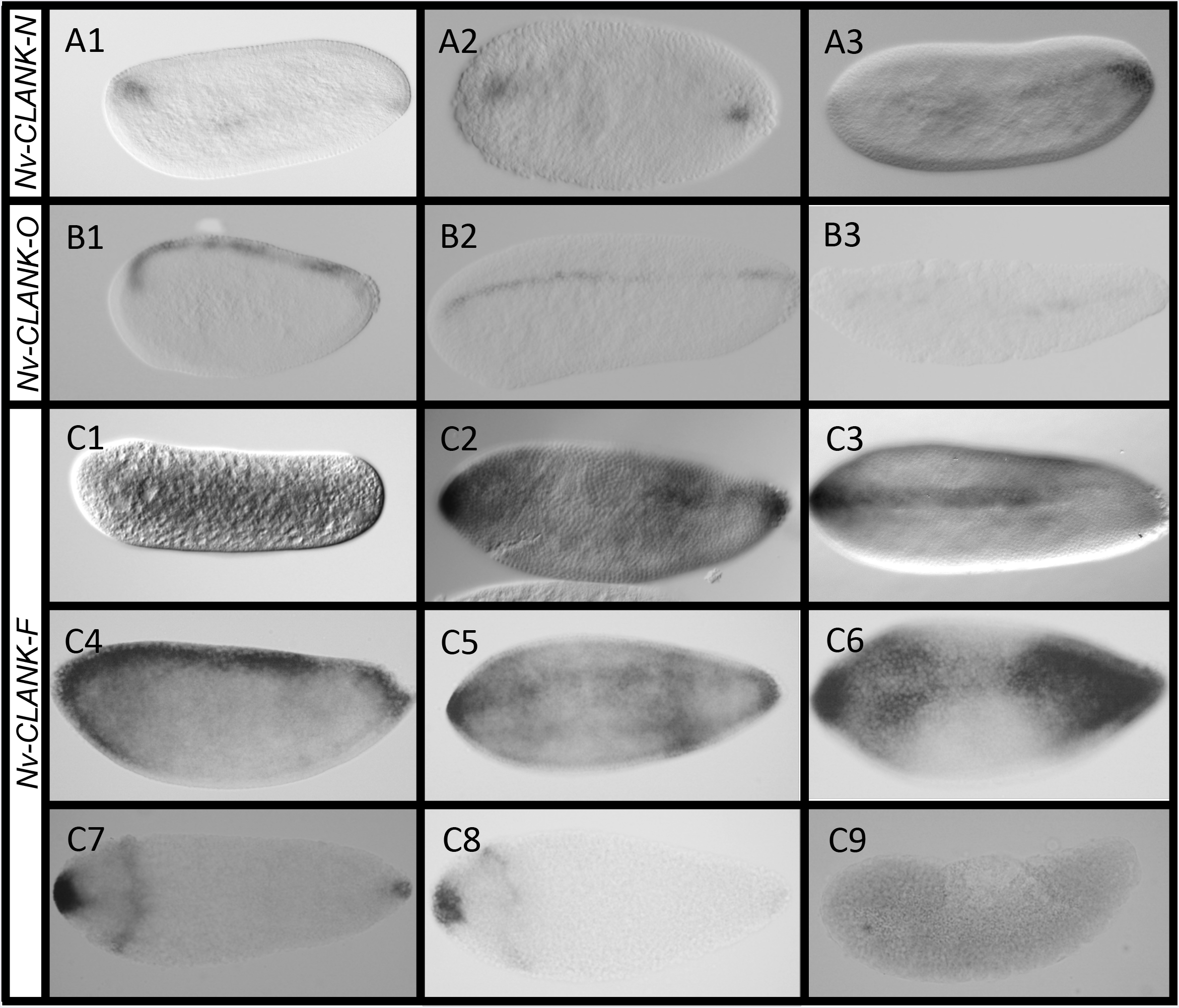
Dorsally expressed CLANKs. A1-A3. Expression of *Nv-CLANK-N* from early to late-blastoderm. B1-B3 Expression of *Nv-CLANK-O* from early-blastoderm through gastrulation. **C1-C9** Expression of *Nv-CLANK-F* from pre-blastoderm through gastrulation. All embryos are oriented with anterior to the left, posterior to the right, dorsal up, and ventral down (except **C7-C8**, bird’s eye dorsal view).

### Ventrally expressed CLANK

*Nv-CLANK-M* is the only transcript in this family with ventral expression. It starts as a narrow stripe in the early blastoderm, much like *Nv-twist* (Fig. 6A1). Also, like *Nv-twi,* it broadens later in development (Fig. 6A2). However, it never takes on a the typical “slug” shape of the presumptive mesoderm and begins to disappear as gastrulation initiates. The pattern of disappearance roughly coincides with regions being covered by the lateral ectoderm (Fig. 6A3).

**Fig 6.**
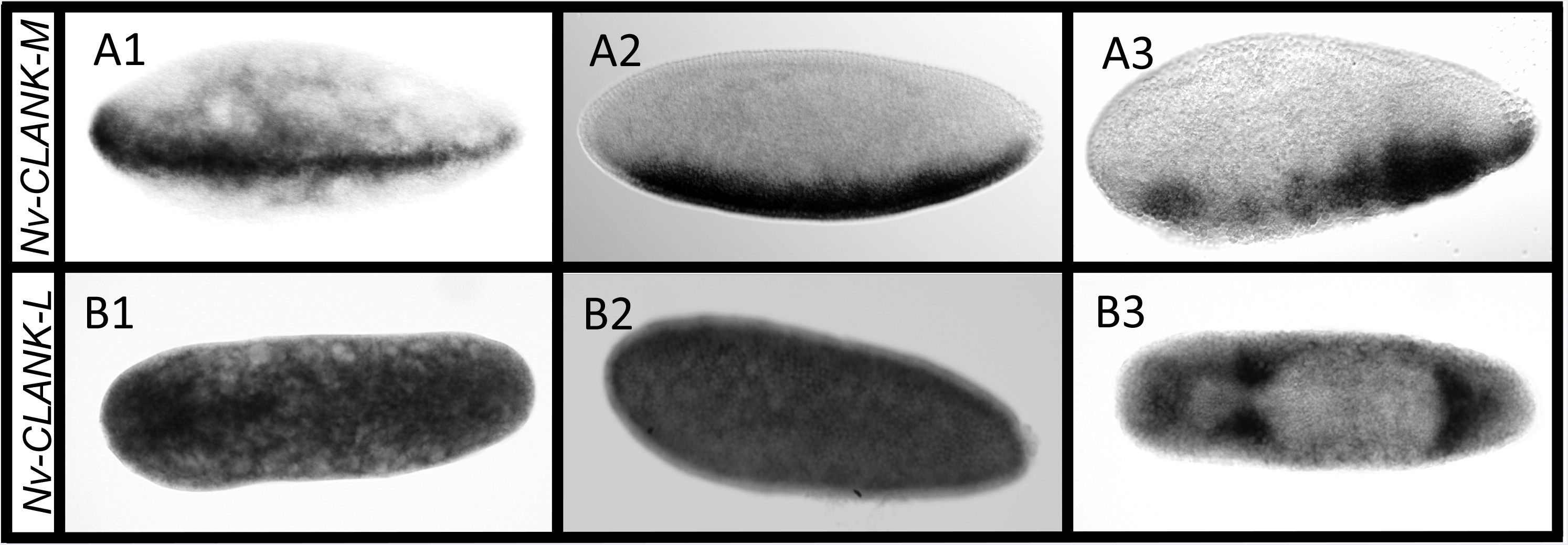
Ventral and other CLANK expression patterns. A1-A3. Expression of *Nv-CLANK-M* from early-blastoderm through gastrulation. **B1-B3** Expression of *Nv-CLANK-L* from pre-blastoderm through gastrulation. All embryos are oriented with anterior to the left, posterior to the right, dorsal up, and ventral down.

### Ubiquitously and postgastrular expressed CLANK

*Nv-CLANK-L* is strongly expressed both maternally (Fig. 6B1) and zygotically (Fig. 6B2-B3). Prior to gastrulation, expression is ubiquitous with the exception of there being no detectable expression within the budding pole cells (Fig. 6B2). During gastrulation there is a moderate level of expression throughout the embryo; however, there are highly elevated levels of expression in the area that will become the head and in regions just anterior and posterior to the extraembryonic material (Fig. 6B3).

### Reduction of CLANK transcripts results in significant increases in embryonic lethality

In order to understand the functional significance of this DV expression of the CLANKs in *Nasonia,* parental RNA interference (pRNAi), where double stranded RNA is injected into female pupae and its effects are examined in the embryos she produces [27], was used to knockdown each of the 11 genes with detectable DV expression patterns. We first analyzed the gross effects of knockdown on embryonic survival to hatching first instar larvae. Average embryonic lethality of the 11 knockdowns ranged from 0.87% to 12.19% of embryos plated (Fig. 7). In all cases the frequency of lethality was larger than in control-injected pupae (0.65%). The difference in lethality was statistically significant (p < 0.05) for 6 of the 11 transcripts tested (Fig. 7.

**Fig 7.**
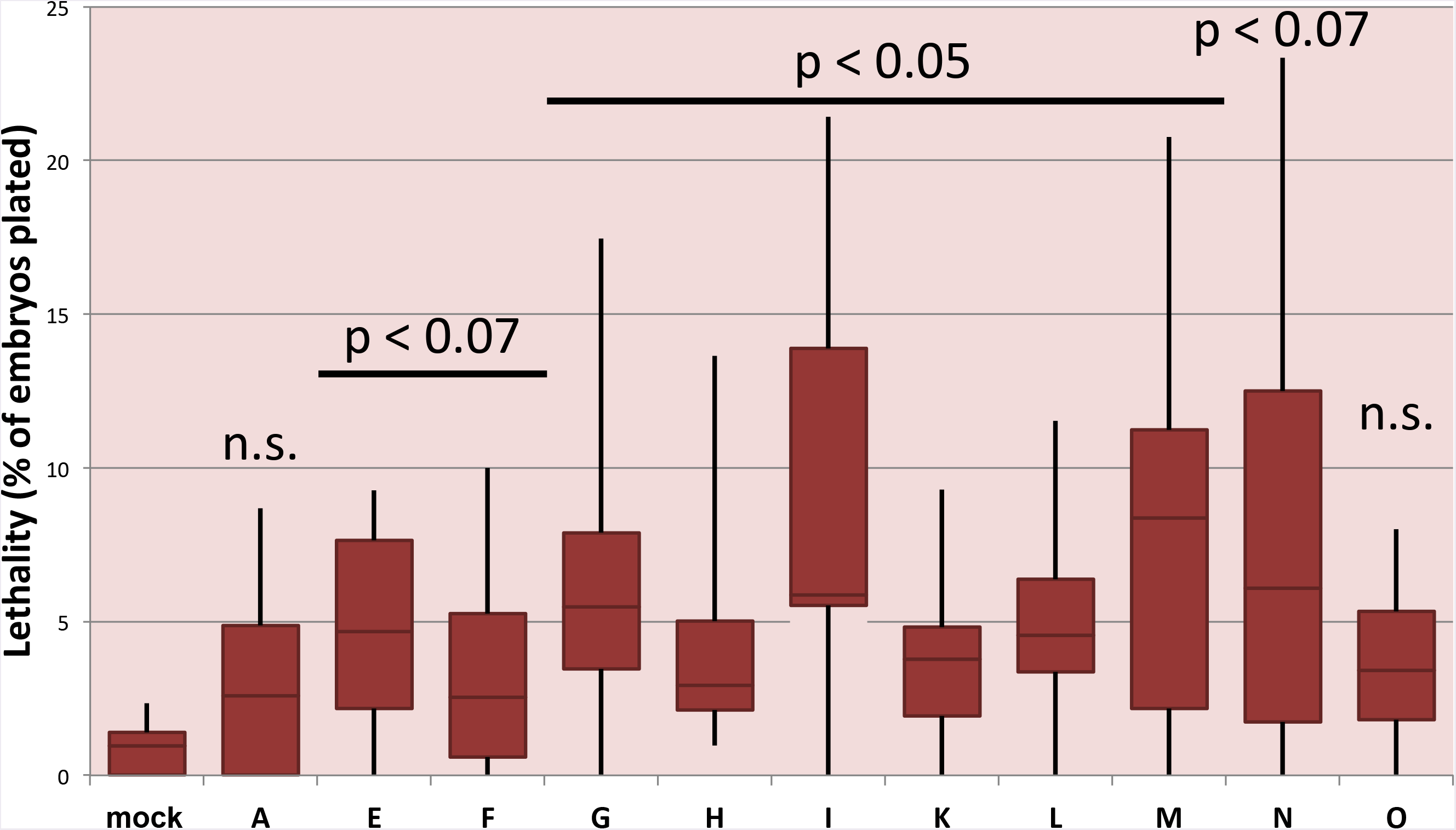
Distribution of pRNAi induced embryonic lethality for each CLANK of interest. Range of embryonic lethality (as a percentage) observed in clutches of embryos from pRNAi knockdown females for each CLANK and mock injected embryos. Error bars represent minimal and maximum values. Horizontal line represents median value. Red box ranges from lower to upper quartile values. t-tests were performed comparing each CLANKs lethality to mock injected lethality. Corresponding p values listed above graph (n.s. = non-significant).

In the course of these experiments focusing on embryonic development, we observed that the pupal injections also had significant and severe effects on successful pupal development for Nv-CLANKs. The effect was particularly strong in *Nv-CLANK-E,* -K, -L where a quarter or less of injected pupae completed metamorphosis, compared the 60% rate for mock injected wasps. (Additional File 7). This suggests that these transcripts may have additional functions in developmental or physiological processes of pupation.

### CLANK transcripts levels are effectively reduced through pRNAi injections

A major caveat of the pRNAi is that the degree by which a transcript is knocked down and the rate at which the system is turned over can vary from gene to gene. In order to test the effectiveness of each designed dsRNA, we quantified the knockdown using qPCR. Ten out of eleven transcripts were reduced to an expression level less than half of that of mock-injected embryos. Their average expression ranged from 7% to 34% of wildtype mRNA expression (Additional File 8). *Nv-CLANK-O* was not as effectively knocked down, but it was still reduced to ~64% of wildtype expression. Expression levels were monitored up to three days post eclosion, we found large variability in the behavior of the dsRNAs (Additional File 8). Some transcripts were immediately reduced, while others required a day to have an observable effect. Additionally, some transcripts were reduced for a number of days, while others quickly regained expression (Additional File 8).

We were encouraged that, despite the incomplete knockdown observed with qPCR, we still saw a significant increase in embryonic lethality. We sought to understand whether this lethality was due to disruptions in patterning in the early embryonic stages, where these CLANKs are expressed. We chose markers of dorsoventral patterning output that were strong, well understood, and represented the embryonic regions most sensitive to patterning disruption. Exactly how the observed disruptions come about is not known and will be the focus of future research.

### Reduction of dorsolaterally expressed CLANK transcripts disrupts patterning specifically on the dorsal side

The *Nasonia* ortholog of *zerknüllt (Nv-zen)* is a well-established marker of the dorsal half of the embryo during normal development [13]. This zygotically expressed transcript is first observed in a broad stripe along the dorsal midline of the early blastoderm (Fig. 8A1, B1). This stripe stretches from the anterior to the posterior pole and is more or less equal in width and intensity throughout its domain. As the blastoderm continues to divide the domain narrows (Fig. 8A2, B2), and eventually retracts from the posterior pole as the blastoderm cellularizes and gastrulation begins (Fig. 8A3, B3). During gastrulation, *Nv-zen* marks the serosa until it begins to migrate and encompass the embryo (Fig. 8A4, B4). Because *Nv-zen* is the most consistently strongly expressed and most well characterized marker on the dorsal side of the embryo, it is an ideal marker to detect disruptions of patterning in this region of the embryo.

**Fig 8.**
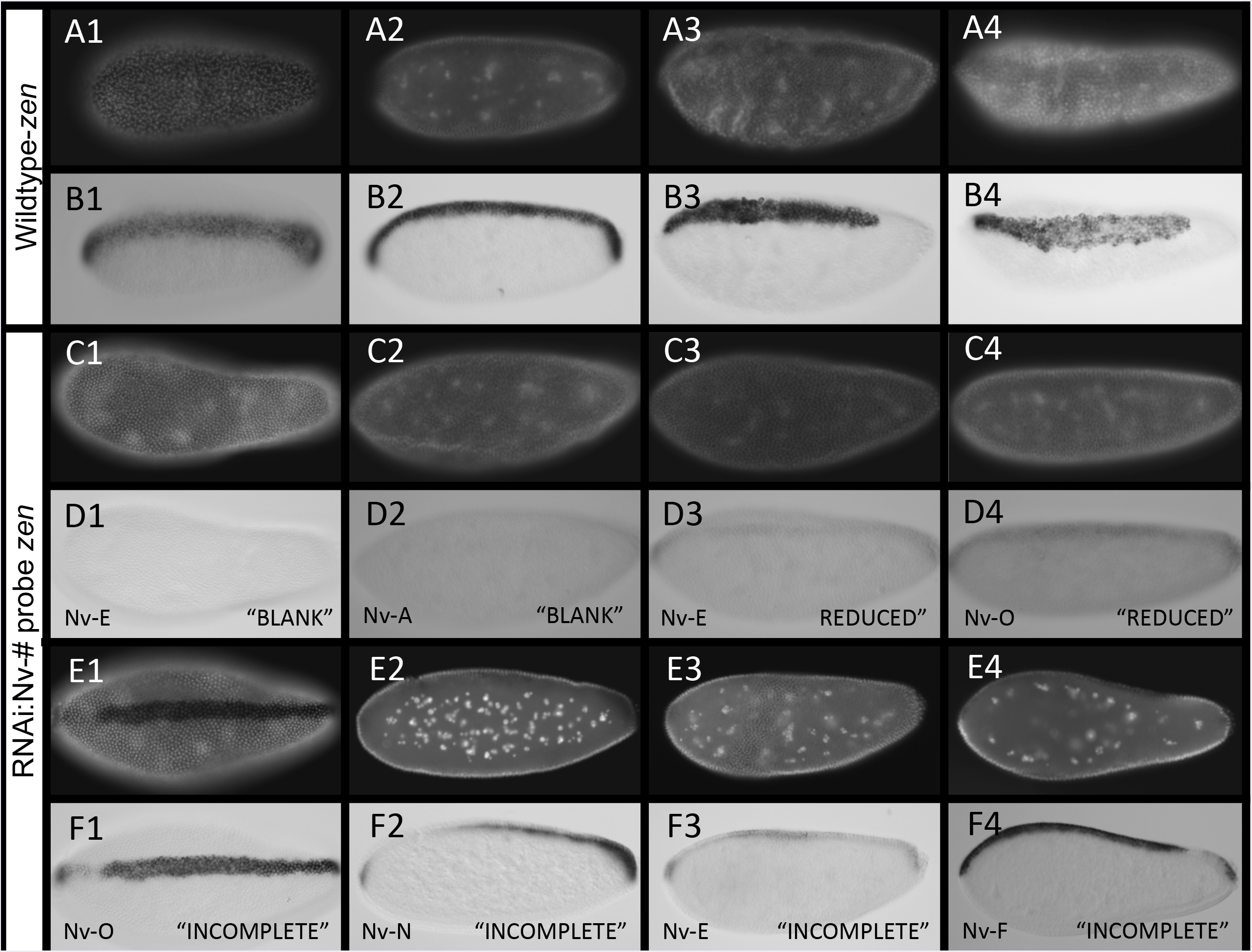
Effects of reducing CLANKs on *Nv-zen* expression. A1-B4. Expression of *Nv-zen* from early-blastoderm through gastrulation in control embryos. **A1-A4** Control embryos stained with DAPI to approximate embryo age. **B1-B4** *In situ* hybridization of control embryos probing for *Nv-zen* expression. Embryos (**B1-B4**) correspond to same embryos in **A1-A4**. **C1-F4** Altered expression of *Nv-zen* following pRNAi of one CLANK transcript (lower left corner) in mid-late blastoderm embryos. **C1-C4**, **E1-E4** Knockdown embryos stained with DAPI to approximate embryo age. **D1-D4**, **F1-F4** *In situ* hybridization of knockdown embryos probing for *Nv-zen* expression (“phenotype” observed, lower right). Embryos correspond to same embryos in **C1-C4**, **E1-E4**. All embryos are oriented with anterior to the left, posterior to the right, dorsal up, and ventral down.

When individual laterally or dorsally expressed CLANK transcripts *(Nv-CLANK-A,* -E, -F, -G, -H, -/, -K, -N, -O) are knocked down, a number of changes in the expression of *Nv-zen* are observed (Fig. 8C1-F4, Additional File 9). First, in all transcripts tested (except *Nv-CLANK-K)* some embryos exhibited reduced levels of *Nv-zen* expression. When visible, the pattern of expression remained unchanged, however, the intensity of the stripe was much lower than control embryos that were processed in the same ISH experiment (Fig. 8D3-4). In other instances, the levels were too low to detect and the embryos appeared to be blank and absent of any *Nv-zen* expression (Fig. 8D1-2). For all of the knockdowns except for *Nv-CLANK-A* and *-K,* the increased frequency of reduced levels of *Nv-zen* expression was statistically significant (p<0.05) (Summarized in Fig. 9).

**Fig 9.**
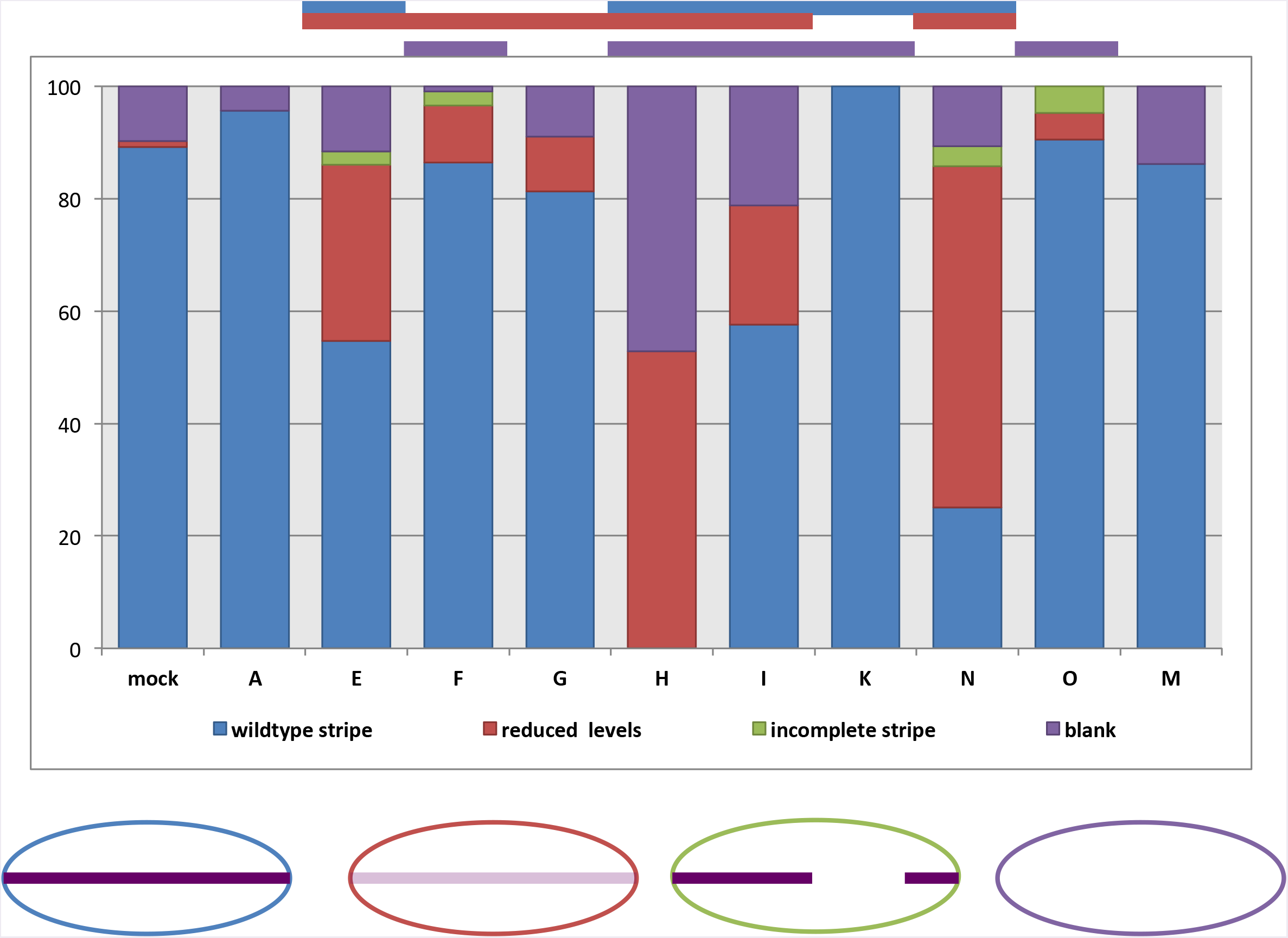
Distribution of pRNAi phenotypes affecting *Nv-zen* expression for each knockdown condition. Percentage of knockdown embryos observed with wildtype *Nv-zen* expression, reduced levels of *Nv-zen* expression, an incomplete or partial dorsal stripe domain of *Nv-zen,* or lacking *Nv-zen* expression completely. Mock-injected embryos were also observed for comparison and to calculate Fisher’s Exact Test to determine if a significant difference (p < 0.05) between the two populations for the given phenotype exists (p-value < 0.05 signified by bar above graph, color corresponds to phenotype with significant difference). Schematic representation of each phenotype is shown below graph.

The second change observed is in the spatial domain of *Nv-zen*. The continuity of the *Nv-zen* expression domain is interrupted in a small proportion of embryos resulting from knockdown of *Nv-CLANK-E, -F, -N,* and *-O*. In mild cases, a small region adjacent to either the anterior or posterior pole is lacking expression (Fig. 8F1, F4), while the proximal pole and all distal regions appear unchanged. In more severe cases, larger regions, up to half the embryo (Fig. 8F2), or multiple regions throughout the embryo (Fig. 8F3) are lacking *Nv-zen* expression. This “incomplete stripe” phenotype was never observed in wildtype embryos. *Nv-CLANK-K* exhibited no abnormalities in *Nv-zen* expression (Summarized in Fig. 9).

To determine whether the spatial patterns of expression of the CLANKs are related to their regions of activity, the ventrally expressed *Nv-CLANK-M* was knocked down and, as would be expected if its function is restricted to its region of expression, the reduction of this ventrally expressed gene had no apparent effect on patterning the dorsal side of the embryo, as all observed embryos appeared phenotypically wildtype, displaying strong dorsal staining of *Nv-zen* (Summarized in Fig. 9).

### Reduction of ventrolateral CLANK transcripts disrupts patterning, morphogenetic movements, and relative timing of embryonic events

Like *zen, twist (Nv-twi)* is a well-established marker of embryonic development in *Nasonia,* but for the ventral region of the embryo [13]. *Nv-twist* is first expressed in a thin stripe along the entire ventral midline of the early blastoderm (Fig. 10A1-B1). As the blastoderm undergoes additional divisions the stripe widens (Fig. 10A2-B2) before retracting at the anterior pole forming a pattern that resembles that of a slug (Fig. 10A3-B3). This slug shape persists through cellularization and into the onset of gastrulation and marks the presumptive mesoderm specifically. The shape is lost once the mesoderm begins to internalize at the anterior end of the domain. This internalization progresses from anterior to posterior (Fig. 10B4) until the entire mesoderm is covered by neuroectoderm.

Ventral and laterally expressed CLANK transcripts *(Nv-CLANK-A,* -E, -G, -H, -/, -K, -M) were knocked down individually, and the expression pattern of *Nv-twi* was observed and characterized with *in situ* hybridization probes in a similar manner as with *Nv-zen* (Fig. 10C1-F4, Additional File 10). The reduction of these transcripts leads to a wide array of phenotypes.

**Fig 10.**
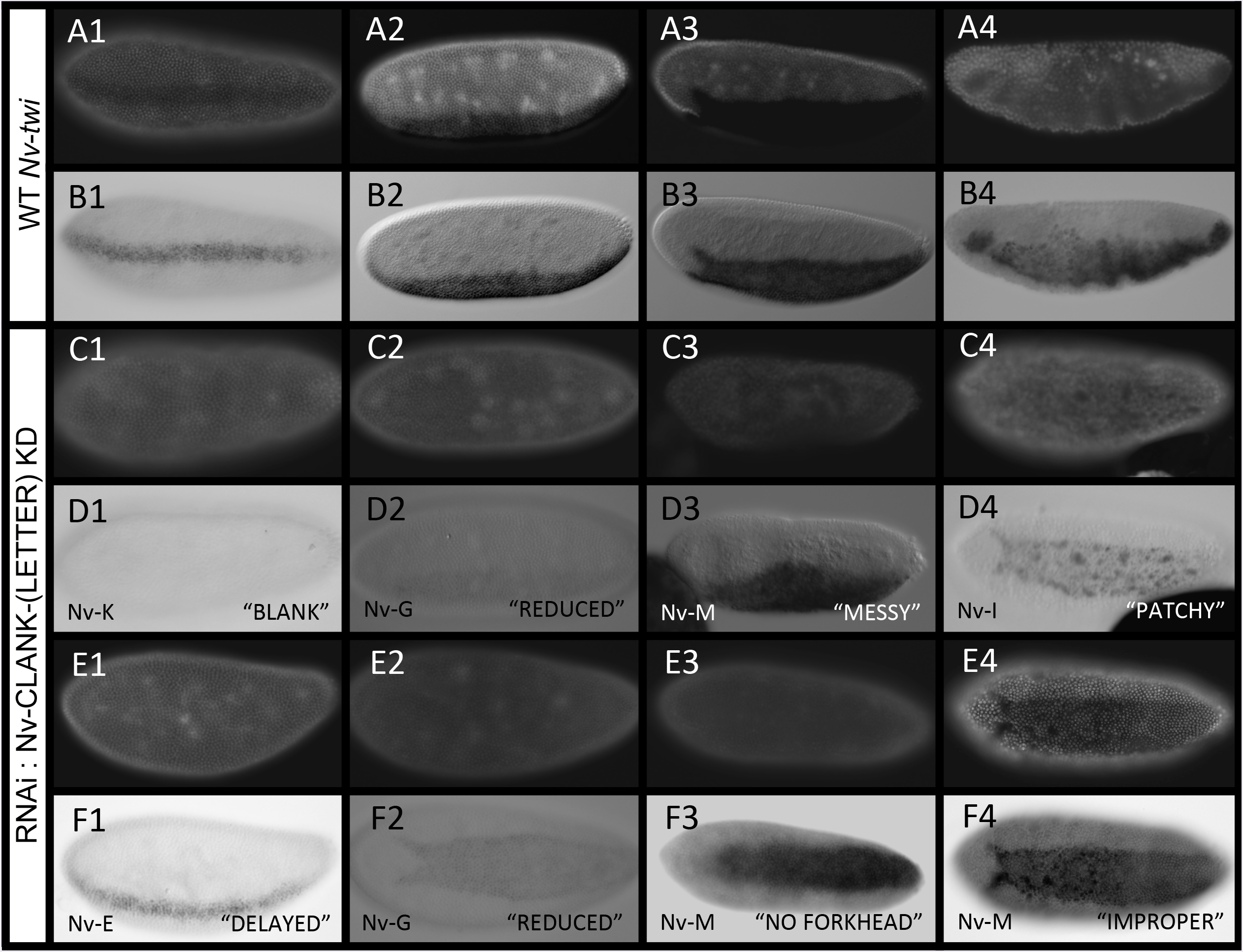
Effects of reducing CLANKs on *Nv-twi* expression. A1-B4. Wildtype expression of *Nv-twi* from early-blastoderm through gastrulation. **A1-A4** Wildtype embryos stained with DAPI to approximate embryo age. **B1-B4** */n situ* hybridization of wildtype embryos probing for *Nv-twi i* expression. Embryos (**B1-B4**) correspond to same embryos in **A1-A4**. **C1-F4** Altered expression of *Nv-twi* following pRNAi of one CLANK transcript (lower left corner) in early-blastoderm through gastrulating embryos. **C1-C4**, **E1-E4** Knockdown embryos stained with DAPI to approximate embryo age. **D1-D4**, F1-F4 *in situ* hybridization of knockdown embryos probing for *Nv-twi* expression (“phenotype” observed, lower right). Embryos correspond to same embryos in **C1-C4**,**E1-E4**. Most embryos are oriented with anterior to the left, posterior to the right, dorsa1139 l up, and ventral down (C4/D4, E2/F2-E4/F4 are birds eye ventral views).

The first group of phenotypes occurs in the early blastoderm when *Nv-twi* expands from a narrow to a wide ventral stripe (Fig. 10A1-B2). The reduction of *Nv-twi* expression is the first phenotype observed at this time point. Structurally these embryos appeared normal, and when present, the spatiotemporal domain of *Nv-twi* is unchanged. The frequency of embryos showing no or lower than normal *Nv-twi* expression (Fig. 10C1-9D2) appeared to be higher in all of the knockdowns except *Nv-CLANK-M,* was statistically significantly different from control for only for *Nv-CLANK-G and -/* (Fig. 11.

**Fig 11.**
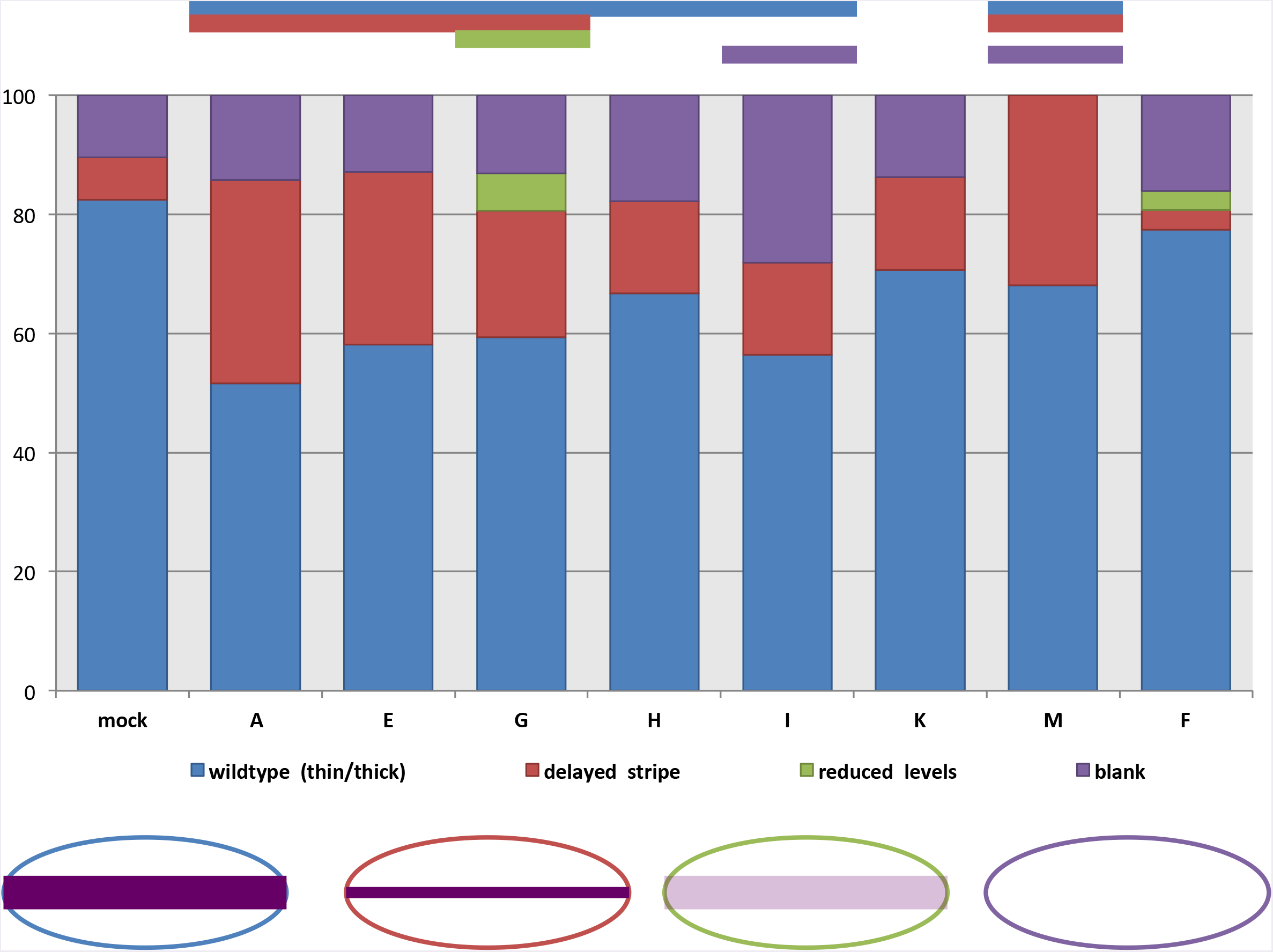
Distribution of pRNAi phenotypes affecting early *Nv-twi* expression for each knockdown condition. Percentage of knockdown embryos observed with wildtype *Nv-twi* expression, a delay in the expansion of *Nv-twi* from a thin to thick ventral stripe, reduced levels of *Nv-twi* expression, or lacking *Nv-twi* expression completely. Mock-injected embryos were also observed for comparison and to calculate Fisher’s Exact Test to determine if a significant difference (p < 0.05) between the two populations for the given phenotype exists (p-value < 0.05 signified by bar above graph, color corresponds to phenotype with significant difference). Schematic representation of each phenotype is shown below graph.

The second phenotype observed in the early blastoderm is a delay in the expansion of *Nv-twi* (Fig. 10F1). The expansion of the *Nv-twi* domain is stereotyped and occurs between nuclear cycle 10 and 11 [13] (compare Fig. 10A1, 10A2, 10E1). This delay phenotype is observed at a frequency higher than in wildtype embryos, after knockdown of all of the lateral/ventral CLANK transcripts, but is only significantly higher for *Nv-CLANK-A,* -E, -G, -M. (Fig. 11.

Effects of CLANK knockdown become more frequent, severe, and varied in the late blastoderm stage, when *Nv-twi* is normally expressed in a ventral “slug” shaped domain (Fig. 10B3). Again, many embryos exhibit reduction in the expression levels of *Nv-twi*. Levels are sometimes reduced completely, as seen after knockdown of *Nv-CLANK-A,* -E, -G, -I, and -K (significantly increased frequency in all but -A, Fig. 12), or to a much lower level than observed in wildtype embryos *(Nv-CLANK-G,* Fig. 10F2, significantly increased frequency, Fig. 12).

**Fig 12.**
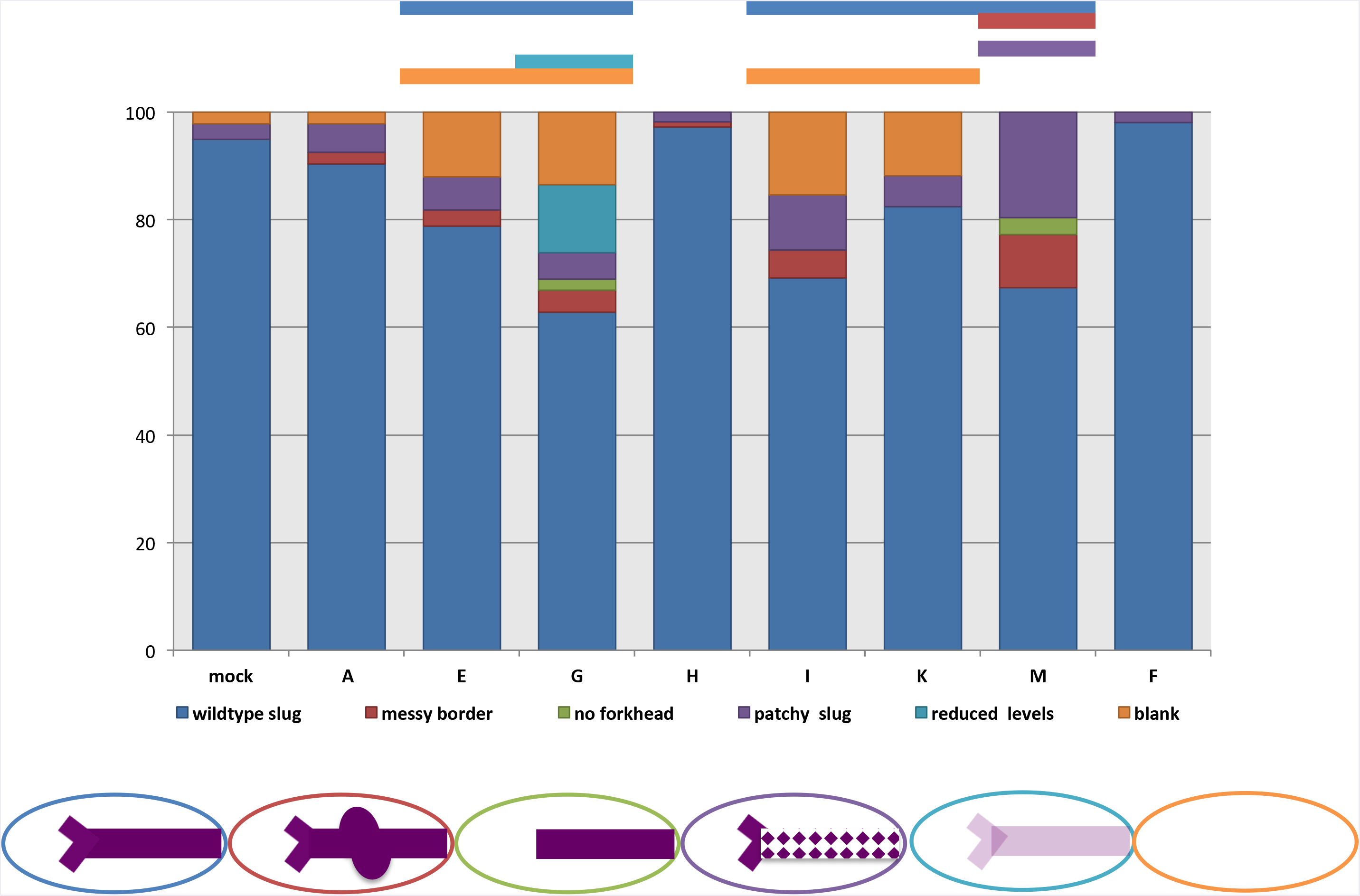
Distribution of pRNAi phenotypes effecting mid-late blastoderm *Nv-twi* expression for each knockdown condition. Percentage of knockdown embryos observed with wildtype *Nv-twi* expression, a messy slug domain border of *Nv-twi,* missing/disrupted slug fork head expression, patchy slug expression, reduced levels of *Nv-twi* expression, or lacking *Nv-twi* expression completely. Mock-injected embryos were also observed for comparison and to calculate Fisher’s Exact Test to determine if a significant difference (p < 0.05) between the two populations for the given phenotype exists (p-value < 0.05 signified by bar above graph, color corresponds to phenotype with significant difference). Schematic representation of each phenotype is shown below graph.

*Nv-CLANK-A,* -E, -G, -H, -I, or -M knock downs all lead to a disruption in the slug shaped domain of *Nv-twi*. Normally this pattern has very sharp, straight lateral borders and a very distinct forking at its anterior end. The sharpness and straightness of the lateral borders are affected at a low, but consistent, frequency after *Nv-CLANK-A, -E, -G, -H, -I,* and -M knockdown (Fig. 10D3). In rarer cases, *Nv-twi* domains where the anterior fork did not resolve are observed (only in *Nv-CLANK-G* and -M, Fig. 10F3). Finally, in some embryos the edges of the *Nv-twi* domain remain unchanged, but within the domain, large patches of cells lack *Nv-twi* expression (Fig. 10D4). The size and number of these patches vary from embryo to embryo within and between knock down conditions.

This “patchy” phenotype was observed after knockdown of all seven ventral/lateral *CLANKs,* and at a much lower frequency in control embryos. However the difference in frequency of this phenotype was significantly only in *Nv-CLANK-M* knockdowns relative to control (Fig. 12. The “messy” border and missing anterior fork phenotypes were never observed in wildtype embryos (Fig. 12.

The last time point where we looked for disruption is during gastrulation. Again, embryos were observed that lack positive staining for *Nv-twi* expression as with the two early stages of development (data not shown); however, this also occurred in wildtype embryos, and only the knockdown of *Nv-CLANK-K* resulted in a frequency of this phenotype significantly higher than what is expected (Fig. 13.

**Fig 13.**
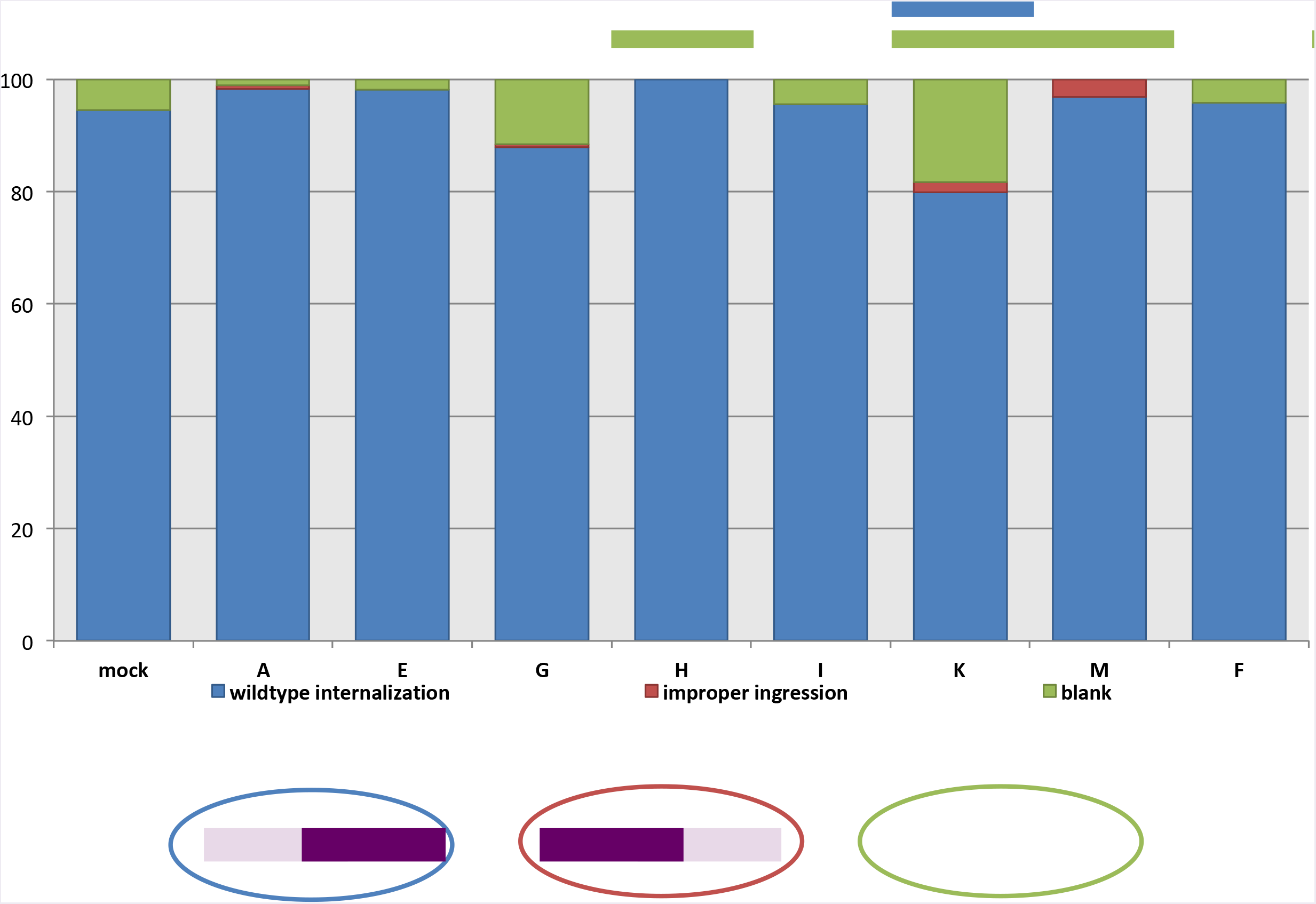
Distribution of pRNAi phenotypes effecting late *Nv-twi* expression for each knockdown condition. Percentage of knockdown embryos observed with wildtype *Nv-twi* expression and mesodermal internalization, improper ingression of the mesoderm, or lacking *Nv-twi* expression completely. Mock-injected embryos were also observed for comparison and to calculate Fisher’s Exact Test to determine if a significant difference (p < 0.05) between the two populations for the given phenotype exists (p-value < 0.05 signified by bar above graph, color corresponds to phenotype with significant difference). Schematic representation of each phenotype is shown below graph.

More interestingly, some knockdowns disrupted the morphogenetic movements of gastrulation. Normally mesoderm internalization proceeds from anterior to posterior in *Nasonia* [13] (Fig. 10B4). In rare instances, the mesoderm was observed internalizing posteriorly to anteriorly (Fig. 10F4) or in a random, disorganized manner (Additional file 10D, Q, Y, Z, EE) when *Nv-CLANK-A,* -G, -K, or -M are knocked down. While never observed in control embryos (or in the myriad normal embryos observed in other experiments), this phenotype occurred at the lowest frequency of all that have been described, and in no condition is the frequency statistically significant (Fig. 13.

To again test whether the spatial expression of the CLANKs is correlated with the location of their phenotypic effects, knockdown embryos from *CLANKs* expressed on the dorsal half of the embryo *(Nv-CLANK-F, -N,* -O) were also examined for changes in *Nv-twi* expression. As expected, the loss (or reduction) of these dorsal transcripts had no effect on patterning of the ventral side of the embryo. All observed embryos appeared phenotypically wildtype, displaying strong *Nv-twi* staining *(Nv-CLANK-F:* Fig. 11-13 *Nv-CLANK-N, -O:* data not shown).

In summary, all of the 11 tested genes showed an increase in embryonic lethality compared to control. We could then show that markers of DV cell fates are disrupted spatially (for both *Nv-twi* and *Nv-zen)* and temporally *(Nv-twi* expansion) by knockdown of different CLANKs. This shows that these novel components of the DV GRN are functionally integrated and are important in producing a stable and reproducible patterning output.

The above results led us to wonder how long these genes have been a part of DV patterning in the wasp lineage to *Nasonia*. Are all of these genes unique recent additions to *Nasonia* DV GRN, or do some of them have a longer history in the wasp lineage?

### Discovery of CLANKs in the wasp *Melittobia digitata*

The second approach to understand the developmental and evolutionary significance of this gain of DV expression in *Nasonia* was to examine the function and expression of CLANKs in other species. This will help to understand how these genes have been functionally integrated into developmental processes.

As we described above, it appears that CLANKs are an ancestral and unique feature of the Superfamily Chalcidoidea. We have chosen to develop *Melittobia digitata,* a representative of the Family Eulophidae (separated from *Nasonia* by about 90 million years of independent evolution [28], as a comparative model. *Melittobia* is attractive because it is easily reared in the lab on the same hosts as *Nasonia,* its mode of embryogenesis is rather similar to *Nasonia,* allowing for more straightforward comparisons of expression patterns, and it adds an important phylogenetic sampling point in understanding the evolution of development within the megadiverse Chalcidoidea.

We sequenced and assembled an embryonic transcriptome from *Melittobia,* and then searched for potential orthologs of *Nasonia* CLANKs within this transcriptome using local BLAST [29]. The sequences of the *potential Melittobia* CLANK homologs are presented in Additional File 5, and were used to generate antisense probes to assess expression patterns (Fig. 14 and in phylogenetic analysis to assess their relationships among themselves and with *Nasonia* CLANKs (Fig. 15.

**Fig 14.**
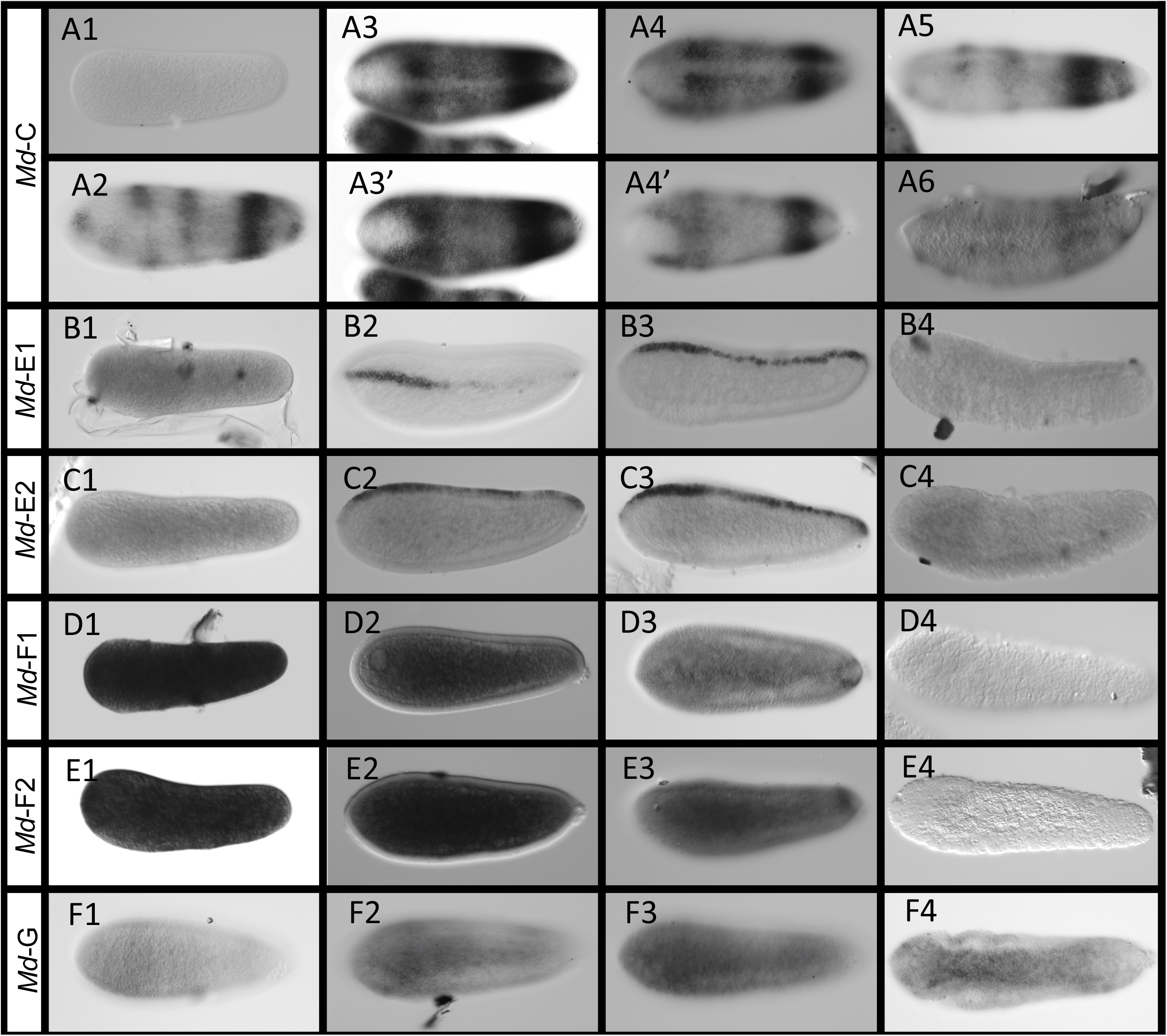
*Melittobia* CLANK candidates with significant expression patterns. A1-A6. Expression of *Md-CLANK-C* from pre-blastoderm through gastrulation. **A3,A4** are dorsal views. **A3**’,**A4**’ are ventral views of the same embryo. **B1-B4** *Md-CLANK-E1* from pre-blastoderm through gastrulation. **C1-C4** *Md-CLANK-E2* from pre-blastoderm through gastrulation. **D1-D4** *Md-CLANK-F1* from pre-blastoderm through gastrulation (**D3** is a bird’s eye, dorsal view). E1-E4 *Md-CLANK-F2* from pre-blastoderm through gastrulation. **F1-F4** *Md-CLANK-G* from early blastoderm to the start of gastrulation. F2-F4 are bird’s eye ventral views. All embryos are oriented with anterior to the left, posterior to the right, dorsal up, and ventral down (unless otherwise noted).

**Fig 15.**
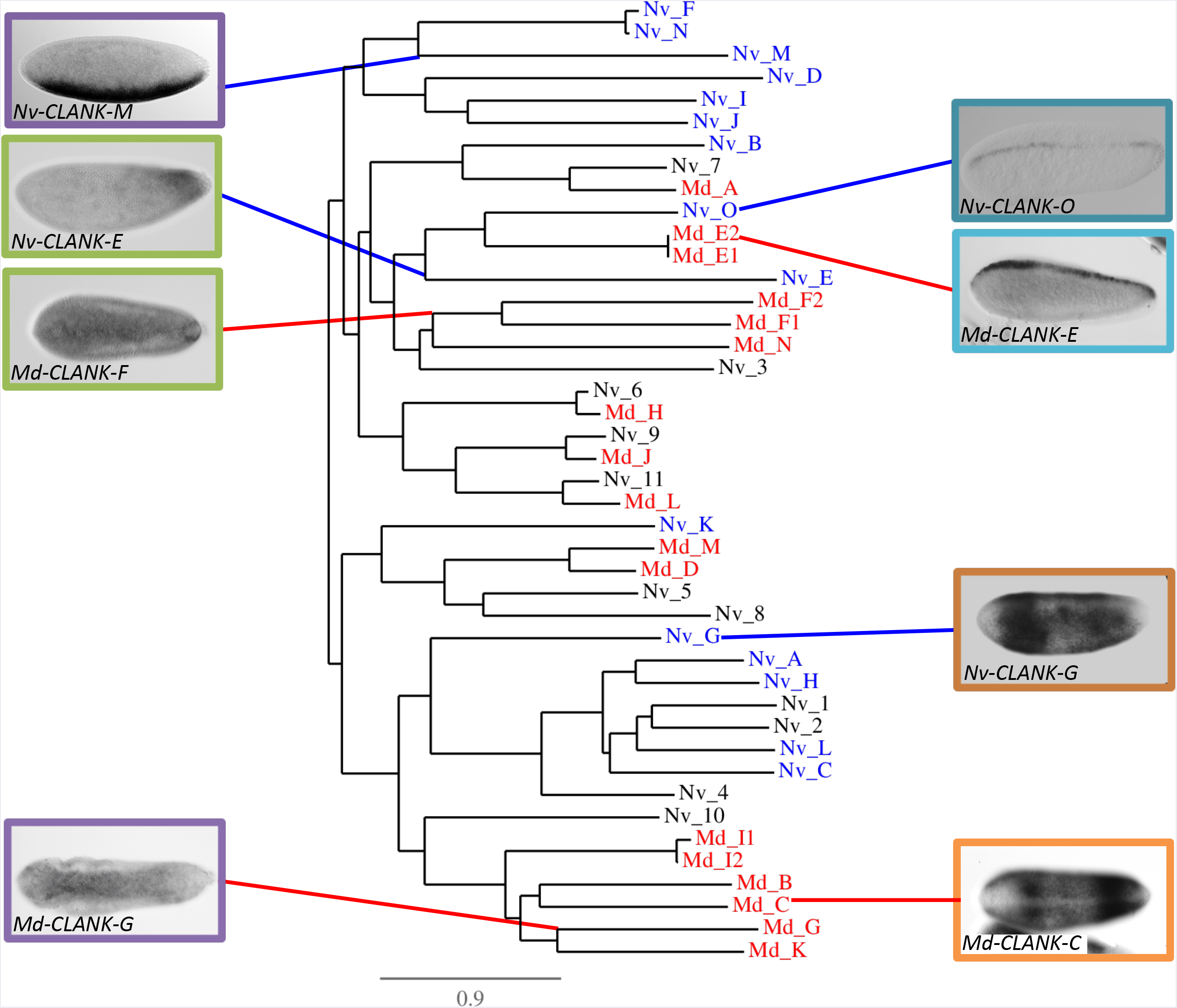
Phylogenetic analysis of *Nasonia* and *Melittobia* CLANK protein families. (**Tree**) Phylogenetic tree of CLANK proteins of interest. Blue = *Nasonia* CLANKs. Red = *Melittobia* CLANK ortholog candidates. Black = *Nasonia* off-target sequences from *Melittobia* reciprocal BLASTs. Branch length is proportional to the number of substitutions per site [25]. (**Inset Images**) Representative images of *Melittobia* orthologs with significant RNA localization and *Nasonia* ortholog with similar pattern. Colored box highlights pairing between wasp orthologs (violet, lime, teal, tangerine). Colored lines point to orthologs phylogenetic branch on tree (Red = *Melittobia,* Blue = *Nasonia*).

### Characterization of *Melittobia* CLANK expression

Since there was only weak evidence for direct orthology of *Melittobia* sequences to the *Nasonia* DV CLANKs, we considered all of the *Melittobia* genes we found to be potential homologs and assessed their expression. Ten of the seventeen Md-CLANKs *(Md-CLANK-A,* -B, -D, -H, -/, -/2, -J, -K, -L, -N) we identified as potential homologs of the DV Nv-CLANKS were not expressed differentially along the DV axis (Additional File 11).

In contrast, *Md-CLANK-C* again has dynamic expression in early and gastrulating embryos (Fig. 14A1-A6). It is absent in pre-blastoderm embryos (Fig. 14A1), then is initially expressed in three bands along the AP axis of the embryo (Fig. 14A2). The strongest and most complete is near the posterior pole. Expression increases in strength and size forming a lateral domain that almost encapsulates the whole embryo. Expression is lacking at the two poles, along the dorsal midline, and along most of the ventral midline (Fig. 14A3-A3’). This lateral domain then retracts into two discrete bands of expression (Fig. 14A4-A4’). The anterior most bands then disappears, leaving just one posterior band (Fig. 14A5). Throughout this retraction the lack of staining at the dorsal midline and poles persists, however, there is staining at the ventral midline in the two bands (compare Fig. 14A3-A4’). During gastrulation this posterior band of expression slowly fades until expression is lacking throughout the embryo (Fig. 14A6). This pattern is quite similar to previous patterns seen in *Nv-CLANK-G* and -H. These genes are in a phylogenetic cluster of *Nasonia* CLANKs that is sister to a cluster of exclusively *Melittobia* CLANKs that contains *Md-CLANK-C* (Fig. 15.

*Md-CLANK-E* and *-E2* also have low levels of ubiquitous expression in early embryos and no expression during gastrulation (Fig. 14B1, B4, C1, C4). However, in blastoderm embryos, both are expressed in a stripe along the dorsal midline. The stripe is dynamic in expression levels and size along the entire AP axis for both *CLANKs*. Expression appears to originate at the anterior pole and fill in in a discontinuous manner until the entire dorsal midline is exhibits expression (Fig. 14B2, B3, C2, C3). Satisfyingly, these two genes cluster phylogenetically with *Nv-CLANK-O* (Fig. 15, which is expressed in an almost identical narrow dorsal stripe (Fig. 5B1-B2). This strongly indicates that the common ancestor of these genes was expressed dorsally and that this pattern has persisted for 90 million years.

Expression of *Md-CLANK-F* and *-F2* is dynamic during blastoderm stages of development. Early pre-blastoderm and blastoderm-staged embryos have light ubiquitous expression (Fig. 14D1, E1). Expression is then increased in the yolk of the syncytial blastoderm (Fig. 14D2, E2), reduced to low levels throughout, and then localized in a small patch of the dorso-posterior of the embryo (Fig. 14D3,E3) before quickly being lost again in cellularized blastoderm stage and gastrulating embryos (Fig.14D4,E4). This pattern has no clear counterpart in the *Nasonia* genes we have examined.

In early blastoderm embryos, *Md-CLANK-G* expression appears to be weak and ubiquitous, with slightly higher expression on the ventral half of the embryo (Fig. 14F1). Expression increases in intensity forming a stripe along the ventral midline, widest in the anterior third of the embryo and narrowing in the posterior third (Fig. 14F2-F3). At the onset of gastrulation, expression is strongest and resembles the characteristic slug shaped domain of *Nv-twist,* before staining is lost completely (Fig. 14F4). This pattern does not develop similarly to the ventrally expressed *Nv-CLANK-M* (Fig. 6A1-A3), and we do not consider it homologous.

The phylogenetic analysis revealed that there has likely been large-scale duplication and divergence (and/or gene conversion) in both wasp lineages (Fig. 15. Most of the *Nasonia* DV CLANKs cluster in two distinct clades on either side of the basal split of this protein tree. Similarly, most of the *Melittobia* proteins cluster together, or with the “off-target” *Nasonia* CLANKs that are not involved in DV patterning (Fig. 15. There are only a handful of cases that indicate clear orthology between a *Nasonia* DV CLANK and a *Melittobia* CLANK. Nv-CLANK-O clusters strongly with Md-CLANK-E1 *and -E2,* while Nv-CLANK-K clusters with Md-CLANK-M and -D. For the others, more complex evolutionary histories, involving ancestral genes that duplicated and diverged multiple times in both lineages after their separation must be considered. Thus, we propose that Nv-CLANK-G, -A, -H, -L, and -C derived the same common ancestral gene as Md-CLANK-B, -C, -G, -I1, -I2, and -K. On the other hand, it is unclear how to relate Nv-CLANK-F, -N, -M, -D, -I, and -J to *Melittobia* counterparts.

Horizontally transferred and duplicating genes have been shown to be subject to complex processes of molecular evolution that make defining ancestry of genes particularly difficult [30], which might explain the difficulty in defining orthology in these relatively closely related species. In addition, potentially missing orthologs may have been misassembled in our transcriptome, and thus not picked up by BLAST analysis. However, the genes we now have in hand are already quite informative as to the functional evolution of this gene family.

## Discussion

In this paper we have shown that a group of genes that came about from multiple rounds of gene duplication and divergence events, potentially following one or more HGT events, are stably and functionally integrated into the embryonic DV patterning GRN of the wasp *Nasonia*. Furthermore, we provide evidence that some of the functionally integrated genes have been participating in developmental processes for a long period, extending back at least 90 million years to the common ancestor of *M. digitata* and *N. vitripennis*. These results raise myriad questions about the origin and fate of horizontally transferred genes, why they are sometimes maintained, and how GRNs change in the course of incorporating these invading genes.

## Incorporation, duplication, and diversification of DV CLANKs

Ankyrin-repeat motifs (ANK) are important for protein-protein interactions and are commonly found in proteins across many species [23, 31]. Sequencing of the *Nasonia* genome uncovered that it contains the largest number of genes coding for ANK proteins of any insect [17]. Among this large number of ANK domain containing are orthologs to genes that are conserved features of insect genomes. However, the vast majority (~170) are orphan genes without clear orthologs in other insects that we have termed CLANKs in this manuscript. A clue to the microbial origin of the CLANKs was the discovery of PRANC domains at the C-termini of some of the proteins. The PRANC domain is found in *Wolbachia,* its bacteriophage, poxviruses and various other bacteria, and its presence strongly indicated HGT from *Wolbachia* into the genome of an ancestor of *Nasonia*. Our results using PSI-BLAST to identify cryptic similarity of the C-termini of CLANKS that do not have annotated PRANC domains to C-termini of *Wolbachia* PRANC and ankyrin domain containing proteins further bolsters this case.

Our observations indicate that the large number of CLANKs in *Nasonia* is the result mostly of duplication and divergence of genes present in the most recent common ancestor of the Chalcidoidea, rather than repeated HGTs within the *Nasonia* lineage after the split among the families. This is based on our observation that the proteins are highly diverged from each other and from any presumed ancestor found in *Wolbachia*. In addition, the presence of introns in almost all of the sequences, their clear integration into the transcriptional regulation milieu of the *Nasonia,* and their dispersal throughout the genome strongly indicate that at least the 15 genes we detected as DV regulated have a long history in wasp genomes, and likely have arisen from duplication and divergence processes.

Apparently more recent HGT events may give clues about the origin and evolution of the CLANKs. For example, in the ant *Pseudomyrmex gracilis* [32], and the bee *Ceratina* [33] we find dozens of Ankyrin domain containing genes that cluster together in phylogenetic analyses, and also cluster tightly with Wolbachia in these same analyses (Additional figures 3 and 4). We cannot exclude that multiple (or no) HGT events occurred in the lineage leading to *Nasonia* to give the full complement of CLANK genes in this wasp, and analyses to determine such facts should be an area of considerable effort in the future.

## Why were the ancestors of the *Nasonia* DV CLANKs maintained?

The chances of a gene horizontally transferred from a prokaryote to a eukaryote to be maintained in a genome is likely to be very low, since not only must it gain the ability to be activated and processed by the eukaryotic transcriptional machinery, but it should also quickly gain a function in its new milieu. If these conditions are not met quickly, random mutations will accumulate and without selection, will eventually destroy protein-coding capacity of the transferred sequence (this is true whether a new gene arises by duplication, HGT, or de novo [34-38].

Since ankyrin domains are protein binding domains [31, 39, 40] and direct protein-protein interaction is thought to be an important pre-cursor for proteins to gain new function [41, 42], ankyrin domain encoding genes like the CLANKs may be predisposed to gain function in new environments. Furthermore, assuming the ancestral CLANK possessed a PRANC domain, it could have a ready-made interaction partner in the form of the Nf-κB homolog Dorsal, which has conserved roles in innate immunity and embryonic patterning throughout insects [43].

## Why were CLANKS integrated into DV patterning?

One of the most surprising results of our previous analysis of DV patterning genes in *Nasonia* was the discovery of so many CLANKs with distinct and unique expression patterns. Their potential as important regulators of the Toll/Dorsal pathway was quite exciting, especially as there are still major open questions about how the Toll/Dorsal pathway interacts with BMP signaling to pattern the *Nasonia* embryo [14]. In Poxviruses, PRANC domain containing genes are known to inhibit the activation of the NF-kB pathway, hijacking the innate immune system within their hosts [44]. Additionally, the PRANC domain has been described as being very similar to F-box domains [45], a domain that is known to induce ubiquitination of IkB, its degradation, and the activation of NFkB [46]. These associations with NFkB/IkB are very interesting because while these proteins function as innate immune responses in most mammals, they have been co-opted to have a function in DV patterning of the embryo of higher insects (Dorsal/Cactus), including *Nasonia*. Therefore, it is possible that these CLANKs were incorporated into the DV pathway because they already had previously established interaction domains with proteins within the pathway (Dorsal/Cactus).

That being said, we have made preliminary observations that indicate that integration of CLANKs is pervasive throughout *Nasonia* development. First, our result reported here, that some CLANK pRNAi knockdowns lead to maternal lethality indicates that some of the DV CLANKs have additional roles in adult organisms. In addition, many of our DV CLANKs are clearly also regulated along the anterior-posterior axis, which might indicate that additional CLANKs might also play important roles in the zygotic GRN patterning this axis. Finally, we have identified additional CLANKs showing maternal mRNA localization to both the anterior and posterior poles of the oocyte/early embryo, indicating roles in establishing AP polarity and specifying germ cells, and CLANKs that are specifically upregulated in either male or female embryos, indicating a role in sex determination (JAL, personal observations). Further functional approaches will be undertaken to assess functional integration of CLANKs into these, and additional GRNs.

## What are the molecular roles of DV CLANKs?

While we propose that interaction with Toll/Dorsal signaling may have been important in the initial integration and stabilization of the CLANKS into the ancestral Chalcidoidean genome,, it is not clear that this interaction has been maintained for the modern *Nasonia* CLANKs. The “incomplete stripe” of *Nv-zen* seen after knockdown of *Nv-CLANK-E, -F, -N* and *-O* is reminiscent of weak knockdowns of BMP components in *Nasonia* ( [14], JAL personal observation), whereas knockdown of Toll has no effect on *Nv-zen*. In addition, the strong reduction of intensity of *Nv-zen* could be a disruption of the BMP signaling, or transcription of target genes downstream of this pathway.

The results where the expansion of the Nv-twist domain is delayed could be ascribed to a disruption of Toll signaling. However, our previous work indicated Toll signaling specifies the initial narrow stripe, while the expansion is mediated by zygotic factors. Similarly, the later disrupted border is likely caused by disruption of interactions of zygotic DV patterning genes, rather than Dorsal itself.

At some level, it is not surprising that the CLANKs we find today have not maintained their hypothetical ancestral interaction partners, given the strong divergence of these genes from each other at the amino acid level, and their 150 million years of evolution. Understanding the molecular interactions that mediate the function of the CLANKs will be a high priority in the coming years.

## When were developmental roles for CLANKs fixed in Chalcidoidea, and how common is recruitment of these genes for developmental processes?

Our curiosity about the evolutionary history of the functional integration of CLANKs into the *Nasonia* DV patterning network led us to establish the wasp *Melittobia* as a satellite model organism. Our results indicate on one hand that there is evidence that CLANKs have been integrated into the DV patterning GRN of some wasps for at least 90 million years (since the divergence of the Pteromalid and Eulophid Families) [27], based on the similarity of expression of two sets of CLANK homologs. On the other hand, we also revealed lineage specific expression patterns for genes in both species, indicating that functional integration of CLANKs is an ongoing process. Sampling wasps in other Families in the Chalcidoidea will be necessary to pinpoint the origin of functional integration of DV CLANKs, and to provide directionality to the changes (i.e., are some functional CLANKs being lost in some lineages, or is the pattern more due to independent gains?). It is likely that an unbiased approach to identify DV regulated genes in *Melittobia* similar to the one we took in *Nasonia* [15], and broad application of this approach in a defined phylogenetic entity would allow for a high resolution characterization of the evolutionary history of this gene family.

## Conclusion

Our knockdowns of DV CLANKs showed that these genes significantly participate in ensuring successful embryogenesis and the formation of viable larvae. Further we showed that a potential role of these genes is to ensure proper establishment of cell fates along the DV axis. However, in neither the patterning nor the viability roles do the CLANKs appear to be absolutely essential. Rather, we propose that CLANKs act to constrain fluctuations in early development, and that their loss can lead to variable fluctuations in patterning which in only rare cases lead to lethality. The source of the fluctuations could be environmental, genetic, or a combination of the two [6, 47, 48]. Our results show that even a very modest contribution to stability of development may lead novel GRN components to be maintained over significant evolutionary time periods. Thus, our results can be extrapolated to any potential novel components of GRNs, whether they originate from HGT, de novo genes, and co-option of existing genes into a new network.

## Methods

### BLASTs and sequence characterization

Our starting sequences were from the *Nasonia* annotation 2.0 [49]. Their NCBI counterparts were found by BLAST against the nt database [50]. These results provided the corresponding NCBI Reference Sequence Accession information for the *Nasonia* 2.0 annotations, which was used for downstream analyses of protein function.

Other similarity searches used blastp and searched the non-redundant protein database and default parameters, except cases where searches were limited to a single species.

### Protein sequence alignment, phylogeny, and conserved domain analysis

Protein sequences corresponding to these fifteen transcripts were then submitted to search for conservation at the amino acid level and to rendered a phylogenetic tree using “One Click Robust Phylogenetic Analysis” (http://phylogeny.lirmm.fr/) [25]. “One Click” parameters were as followed: Data& Settings (Gblocks not used), MUSCLE Alignment (-SeqType Protein), PhyML Phylogeny (Substitution model: WAG), TreeDyn Tree Rendering (Reroot using mid-point rooting, Branch annotation: Branch support values). Finally, in order to gain family, domain and repeat information for each transcript, we analyzed each protein using the Interpro Protein sequence analysis & classification software (InterProScan5) (https://www.ebi.ac.uk/interpro/") [26].

Phylogenetic analysis of the top 100 BLAST hits to each CLANK was performed at http://www.trex.uqam.ca [51]. Alignments were performed using MUSCLE [52], alignments were manipulated for correct file type using AliView [53], relationships were inferred with RAxML [54] and trees were drawn and edited in FigTree: (http://tree.bio.ed.ac.uk/software/figtree/"). Default parameters were used.

PSI_BLAST was performed using sequence downstream of annotated ankyrin domains in *Nasonia* CLANKs. 2 iterations were performed for each gene, except CLANK-L, which required 4 iterations to find sequences outside of *Nasonia* and *Trichomalopsis*. Only sequences above threshold were used to seed the next iteration. All aligning sequences were downloaded as FASTA files. These were aligned all to eachother using MUSCLE implemented at T-rex as above, and the resulting alignments were used for phylogenetic analysis as described above.

### RNA interference, screening, and embryo collection

Yellow AsymCx (wild-type, cured of *Wolbachia)* pupa were injected with dsRNA (~1μg/mL in water) designed against each of the transcripts with significant DV expression as described in [27]. Injected pupae were allowed to eclose and lay eggs. Overnight egg lays were collected and plated onto 1% PBS agar plates. Embryos were aged 24 hours at 28°C and then screened for embryonic lethality. Mock, water-injected embryos were also collected, plated, and screened as a control. Nearly all water-injected embryos are predicted to hatch within 24 hours and develop into crawling, feeding, larva. A small number of embryos will fail to hatch, and instead development will arrest at the embryonic stage. We define this failure to hatch as embryonic lethality. If the transcript we knockdown is predicted to be vital for embryonic development, we predict that a higher percent of injected embryos will exhibit this embryonic lethal phenotype compared to mock-injected embryos.

Average embryonic lethality was calculated and plotted as a bar graph for each condition. Standard deviation was used to calculate standard error, and T-tests (p < 0.05) were used to test for significance. Box plots were also created to provide visualization of the distribution of observed lethality within each conditional population.

Timed egg lays were also conducted to collect 3-7 hours (at 28°C) embryos from each knockdown condition. This time span corresponds to the developmental stages we know these transcripts are differentially expressed (the penultimate syncytial division through the beginning of gastrulation) Embryos were fixed and then processed for in situ hybridization or qPCR.

### Characterization of RNA localization *(in situ* hybridization)

*in situ* hybridization was performed using standard protocols [13, 15] on 0-24 hour, wildtype, AsymCX embryos in order to characterize normal expression patterns of each CLANK transcript during embryogenesis (specific details available upon request). Embryos were imaged at 20X magnification on Zeiss widefield, compound epifluorescent microscope.

For knockdown experiments, *in situ* protocols were repeated on the 3-7 hours (28°C) knockdown embryos and on 3-7 hour mock-injected embryos. Anti-sense probes were generated from primers specific to *Nv-twi* and *Nv-zen*. Knockdown phenotypes were described based on their divergence from mock-injected expression of *Nv-twi* and *Nv-zen,* and their frequency of occurrence was calculated and compared to mock-injected phenotype frequencies. Raw frequency counts were converted to percentages (out of 100) and Fisher’s Exact Test was used to determine if a given phenotypic frequency observed in knockdown embryos was significantly different from the frequency of that phenotype in mock-injected embryos (p <0.05).

### Qualitative Polymerase Chain Reaction (qPCR)

RNA was isolated from 3-7 hours (28°C) embryos using standard TRIzol-based protocols (Ambion 15596018) and converted into cDNA using the Protoscript First Strand cDNA synthesis kit (NEB 63001), controlling for total RNA input. Two cDNA replicates were synthesized per condition. cDNA was synthesized in this manner for each condition for three consecutive days post eclosure of the injected wasps.

To assess knockdown, we performed qPCR on knockdown embryos in parallel with mock-treated embryos. These were carried out using primers specific to the transcript of interest while using the housekeeping gene, *rp49,* as a control. 20μL per well PCR reactions were assembled using the PowerUp SYBR Green Master Mix (Applied Biosystems: A25742)(2X MM, 800nM of each primer, cDNA, RFH2O). We performed the reactions in triplicate using the following parameters: (50°C for 2’, 95°C for 2’, 40 cycles of (95°C for 15 sec, 60°C for 60 sec, plate read, 72°C for 60 sec, plate read), 95°C for 2’, gradient 60°C➔95°C (0.2°C for 1 sec).

Average C_T_ was calculated by combining triplicates from both cDNA replicates for each condition. Knockdown average C_T_ ‘s were then normalized to mock injected RP49 levels. Knockdown Delta C_T_ ‘s were calculated and expressed as a relative expression (percentage of wildtype expression). Relative expression was calculated for each condition per day (up to three days) and an average relative expression was calculated over the three-day span. Standard deviation was used to calculate standard error, and T-test (p < 0.05) were used to test for significance.

## *Melittobia* sequences

Total mRNA (1 ug) libraries were created from various developmental time points

(ovaries, pre-, early-, late-blastoderm embryos, male-, female-yellow pupa) of the wasp *Melittobia digitata* using the NEBNext Ultra Directional RNA Library Prep Kit for Illumina (NEB #E7420) in conjunction with NEBNext Poly(A) mRNA Magnetic Isolation Module (NEB #E7490). Libraries were validated and quantified before being pooled and sequenced on an Illumina HiSeq 2000 sequencer with a 100 bp paired-end protocol. Sequences were de novo assembled using Trinity on a Galaxy Portal. (Currently in preparation for publication, specific details available upon request). *Melittobia* RNAseq data is available in the BioSample database under accession numbers [SAMN08361226, SAMN08361227, SAMN08361228].

## *Melittobia* ortholog discovery

We used a de novo assembled, unannotated embryonic *Melittobia digitata* Transcriptome as a database for local tBLASTn (Nv-protein → Md-mRNA) within the Geneious program (https://www.geneious.com/) [29] to search for orthologs to the *Nasonia CLANK* genes. A reciprocal BLAST was done on the top hits. If the reciprocal BLAST resulted in the *Nasonia* sequence that was input into the query being returned as the top hit, the hit was considered a strong ortholog candidate. If it did not correctly BLAST back to the input sequence, the off-target *Nasonia* protein sequence returned was collected to be input in to downstream phylogenetic analysis. *Melitobia* transcript sequences were then translated into amino acid sequences via an online Translate tool (ExPASy,https://web.expasy.org/translate/). Longest ORFs sequences were confirmed to align with tBLASTn hit sequences, and then were collected for phylogenetic analysis.

Input *Nasonia,* off-target *Nasonia,* and all potential *Melittobia* ortholog protein sequences were then submitted to “One Click Robust Phylogenetic Analysis” (http://phylogeny.lirmm.fr/") [25] and trees were rendered relating each chalcid species orthologs to the *Nasonia* sequences as described earlier in our methods.

## *Melittobia in situ* hybridization

Embryos collection, processing, and *in situ* protocol were developed and performed in a manner similar to *Nasonia* protocols with minor modifications. (Currently in preparation for publication, specific details available upon request).

## List of Abbreviations

ANK: Ankyrin-repeat motifs
CLANKs: Chalcidoidea Lineage specific ANKyrin domain encoding genes
DV: dorsoventral
GRN: Gene regulatory networks
HGT: horizontal gene transfer
*htl*: heartless
ISH: *in situ* hybridization
PRANC: Pox proteins Repeats of ANkyrin, C-terminal domain
pRNAi: parental RNA interference
qPCR: Quantitative PCR
sna: snail
twi: twist
zen: zerknüllt
zfh: zinc finger homeodomain

## Declarations

### Ethics approval and consent to participate

Not applicable

### Consent for publication

Not applicable

### Availability of data and materials

*Melittobia* RNAseq data is available in the BioSample database under accession numbers [SAMN08361226, SAMN08361227, SAMN08361228], the SRA database under accession number SRP129036, and the BioProject database under accession number PRJNA429828.

## Authors’ contributions

DP provided assistance in experimental design and performed all the experiments. JAL designed the experiments and supervised students. All authors contributed to writing the manuscript and approved the final manuscript.

## Acknowledgements

We would like to like to acknowledge the DNA Services Facility at UIC for assistance in transcriptome sequencing. We would like to thank Sarah Bordenstein and an anonymous reviewer for helpful suggestions and discussion on this manuscript.

## Additional Files

### Additional File 1. (.xlsx) *D.melanogaster* and *A.mellifera* BLASTp hits for each *Nasonia* novel ankyrin-repeat containing transcript

Each of the 15 novel *Nasonia* transcripts was entered into a BLASTp query to search for homologous proteins in *Drosophila* and *Apis*. The top 100 hits were collected (duplicate sequences were removed) for each *Nasonia* transcript (individual sheets within workbook). Each row represents a distinct BLASTp hit and contains the hit’s description (column A), BLAST e-value (column B), and NCBI accession number (column C).

### Additional File 2. (.xlsx) Top *D.melanogaster* and *A.mellifera* BLASTp novel ankyrin-repeat hits

All sequences from Additional file 1 were pooled (duplicate sequences removed) and entered as a reciprocal BLASTp query to see if they were strongly homologous to the *Nasonia* novel ankyrin-repeat containing transcripts. Each row represents a distinct BLASTp hit from Additioanl file 1, and again contains the hit’s description (column A, Yellow highlights *Apis* sequences. *Drosophila* sequences are left in white), BLAST e-value (column B), and NCBI accession number (column C) as well as the description of the *Nasonia* protein it reciprocal BLASTs back to (column E), BLAST e-value (column F), and NCBI accession number (column G).

### Additional File 3. (.docx)

Phylogenetic Analyses of the top 100 blast hits for each of the 15 Nv-CLANKs analyzed in this paper. CLANK sequences are in red, and bacterial and viral sequences are in green. Clades containing many Trichomonas ankyrin domain encoding genes have been collapsed.

### Additional File 4. (.pdf) Phylogenetic analysis of Psi-BLAST of the C-termini of Nv-CLANKs

CLANK sequences are in red, and bacterial and viral sequences are in green. Large clades containing sequences exclusively from a single species have been collapsed. See label for panel number and CLANK used as query to generate the tree.

### Additional File 5. (.xlsx) Relationships between our sequence nomenclature and gene identification numbers in different annotations. Sheet 1. Reference sequences and names corresponding to each novel ankyrin-repeat containing *Nasonia* transcripts

“Working Code”, “Transcript” number from annotation 2.0 of the *N. vitripennis* genome, and corresponding “Gene Symbol”, “NCBI Reference Sequence” (mRNA), and “NCBI Reference Sequence” (protein) for each transcript of interest. Alternatively, spliced transcripts and proteins are listed as a second row for a given “Transcript.” **Sheet 2. CLANK ortholog candidate sequences in *Melittobia*.** Column A: Working code used throughout paper to identify sequences. Column B: *Melittobia* CLANK sequence accession numbers from de novo embryonic transcriptome (in prep). **Sheet 3. *Nasonia* off-target hits from *Melittobia* CLANK ortholog BLAST**. Column A: Working code used throughout paper to identify sequences. Column B: *Nasonia* sequence accession numbers from annotation 2.0 of the *N. vitripennis* genome. Column C: *Nasonia* NCBI Reference Sequences. Column D: *Nasonia* NCBI Reference Sequences predicted gene name. **Sheet 4. *Melittobia* primer sequences for *in situ* hybridization probes**. Column A: Primer name. Column B: Primer sequence. Primers were designed using Primer3 v.0.4.0 (http://primer3.ut.ee") and synthesized by Integrated DNA Technologies (IDT, http://www.idtdna.com/Site/Order/oligoentry").

### Additional File 6. (.tiff) CLANKs lacking differential expression patterns. A1-D3

Expression of *Nv-CLANK-B, -C,* -D, *-J* from pre-blastoderm through gastrulation. All embryos are oriented with anterior to the left, posterior to the right, dorsal up, and ventral down.

### Additional file 7. (.tiff) Distribution of pRNAi survival rate for each CLANK of interest

Range of pupal survival and eclosure (as a percentage) observed in pRNAi knockdown females for each CLANK and mock injection. Error bars represent minimal and maximum values. Horizontal line represents median value. Red box ranges from lower to upper quartile values.

### Additional File 8. (.tiff) Relative embryonic expression of CLANK transcripts over time following pRNAi

cDNA was generated from aged (3-7 h, 28°C) embryos, collected from pRNAi injected females for up to three days post eclosure. mRNA expression levels of the knockdown transcript were monitored via qPCR. Relative expression compared to mock injected embryos (as a percentage out of 100) was calculated and plotted after reactions were normalized via *Nv-rp49* expression. Expression values are an average of biological and technical replicates.

### Additional File 9. (.tiff) Effects of reducing *CLANKs* on *Nv-zen* expression. A-U’

Altered expression of *Nv-zen* following pRNAi of a *CLANK* of interest in mid-late blastoderm embryos. **A-U** Knockdown embryos stained with DAPI to approximate embryo age. **A’-U**’ */n situ* hybridization of knockdown embryos probing for *Nv-zen* expression. Embryos correspond to same embryos in **A-U**. All embryos are oriented with anterior to the left, posterior to the right, dorsal up, and ventral down. **A-B**’ *Nv-CLANK-A* pRNAi embryos.**C-D**’ *Nv-CLANK-G* pRNAi embryos. **E-G**’ *Nv-CLANK-E* pRNAi embryos. **H-K**’ *Nv-CLANK-F* pRNAi embryos. **L-M**’ *Nv-CLANK-H* pRNAi embryos. **N-O**’ *Nv-CLANK-/* pRNAi embryos. **P-R**’ *Nv-CLANK-N* pRNAi embryos. **S-U**’ *Nv-CLANK-O* pRNAi embryos. Descriptive term of phenotype observed in bottom right corner of *in situ* images.

### Additional File 10. (.tiff) Effects of reducing *CLANKs* on *Nv-twi* expression. A-EE’

Altered expression of *Nv-twi* following pRNAi of a *CLANK* of interest in mid blastoderm to gastrulating embryos. **A-EE** Knockdown embryos stained with DAPI to approximate embryo age. **A’-EE**’ */n situ* hybridization of knockdown embryos probing for *Nv-twi* expression. Embryos correspond to same embryos in **A-EE**. All embryos are oriented with anterior to the left, posterior to the right, dorsal up, and ventral down (unless otherwise noted). **A-D**’ *Nv-CLANK-A* pRNAi embryos (C/C’, D/D’ ventral views). **E-F**’ *Nv-CLANK-E* pRNAi embryos. **G-I**’ *Nv-CLANK-H* pRNAi embryos (H/H’ ventral views). **J-Q**’ *Nv-CLANK-G* pRNAi embryos (K-N’ ventral views). **R-U**’ *Nv-CLANK-/* pRNAi embryos (S/S’, U/U’ ventral views). **V-Z**’ *Nv-CLANK-K* pRNAi embryos. **AA-EE**’ *Nv-CLANK-M* pRNAi embryos (AA-EE’ ventral views). Descriptive term of phenotype observed in bottom right corner of *in situ* images.

### Additional File 11. (.tiff) *Melittobia* CLANK candidates lacking differential expression patterns. A1-J3

Expression of *Md-CLANK-A,* -B, *-D, -H, -/1, -/2,* -J, *-K, -L,* and *-N* from pre-blastoderm through gastrulation. All embryos are oriented with anterior to the left, posterior to the right, dorsal up, and ventral down (except B3, bird’s eye dorsal view).

